# Structural elements of cyanobacterial co-factor-independent phosphoglycerate mutase that mediate regulation by PirC

**DOI:** 10.1101/2024.10.28.619893

**Authors:** Tim Orthwein, Janette T. Alford, Nathalie S. Becker, Phillipp Fink, Karl Forchhammer

**Affiliations:** Interfaculty Institute of Microbiology and Infection Medicine Tuebingen, University of Tuebingen, Germany

## Abstract

The 2,3-bisphosphoglycerate-independent phosphoglycerate mutase (iPGAM) has been identified as a crucial regulating key point in the carbon storage metabolism of cyanobacteria. Upon nitrogen starvation, the iPGAM is inhibited by the P_II_-interacting regulator PirC, released from its interaction partner P_II_ due to elevated 2-oxoglutarate levels. *In-silico* analysis of 338 different iPGAMs revealed a deep-rooted distinctive evolution of iPGAMs in cyanobacteria. Remarkably, cyanobacterial iPGAMs possess a unique loop structure and an extended C-terminus. Our analysis suggests that iPGAM forms a complex with three individual PirC monomers. Complex affinity is affected by the unique loop and the C-terminal structural elements. A C-terminal truncated enzyme showed loss of control by PirC and two-fold increased enzymatic activity compared to the iPGAM-WT. By contrast, deleting the loop structure drastically reduced the activity of this variant. By replacing the WT iPGAM in *Synechocystis* with different iPGAM variants, in which these structural elements were deleted, it became apparent that deletion of the C-terminal element showed a similar overproduction of polyhydroxybutyrate as deletion of the iPGAM-regulator PirC. However, in contrast to the latter, these strains showed higher over-all biomass accumulation, making them a better chassis for a production strain for PHB or other valuable substances than the PirC-deficient mutant. These findings significantly contribute to our understanding of the metabolic pathways in cyanobacteria and open up new avenues for further research in this field, inspiring future investigations and discoveries.

**Importance:** The primordial cyanobacteria were responsible for developing oxygenic photosynthesis early in evolution. Through endosymbiosis, they further evolved into the chloroplasts found in the plant kingdom. Many metabolic pathways within chloroplasts originated from cyanobacteria. However, differences emerged during their long separate evolution, providing insights into the endosymbiotic process. In the metabolic pathways involving fixed CO_2_, the co-factor-independent phosphoglycerate mutase (iPGAM) plays a crucial role by directing the first CO_2_ fixation product, 3-phosphoglycerate, towards critical anabolic pathways. Our findings reveal a distinct evolution of iPGAM within oxygenic photosynthetic organisms. We have identified two specific segments in cyanobacterial iPGAMs that tightly control the cellular carbon/nitrogen state through a specific protein interactor (PiC). This understanding of iPGAM has allowed us to engineer cyanobacterial strains with altered carbon fluxes. Since cyanobacteria can directly convert CO_2_ into valuable products, our results demonstrate a novel approach for developing a chassis for biotechnical use.

## Introduction

Phosphoglycerate mutases (PGAM) are enzymes present in almost every living organism. They connect the major sugar metabolic routes, the Embden–Meyerhof–Parnas pathway (EMP), the Entner-Doudoroff pathway (ED), the oxidative pentose-phosphate pathway (OPP) and its reductive pendant, the Calvin-Benson-Bassham cycle (CBB), with the reactions of lower glycolysis. At this metabolic lynchpin, PGAM converts 3-phosphoglycerate (3-PGA) into 2-phos-phoglycerate (2-PGA), allowing carbon flux into the lower glycolysis and thereby into many pathways of anabolic metabolism, such as synthesis of fatty acids or amino acids. There are two types of PGAM enzymes, which are affiliated with the same superfamily of alkaline phosphates (1): first, the 2,3-bisphosphoglycerate (2,3-BPG) dependent (dPGAM) and second, the 2,3-BPG independent PGAM (iPGAM) (2, 3, 4). Other than dPGAM, iPGAM does not need activation by 2,3-BPG. iPGAMs exist in all plants, algae, some invertebrates, and fungi and are widespread in bacteria (2). In these enzymes, the reversible transfer of the phosphate of the glycerate core from the C3 to the C2 position is achieved in a two-step reaction. 3-PGA first phosphorylates the active site seryl-residue (phosphatase activity, phosphate sub-domain) followed by reorientation of the substrate and re-phosphorylation of the C2-oxygen by phosphotransferase activity (2). Two manganese ions coordinate the reaction, which makes this reaction highly pH-dependent (3). Species of the *Bacillota* (*Firmicutes*) phylum use this effect to regulate iPGAMs, which play a crucial role in endospore formation (4). In analogy to the *Firmicutes*, iPGAMs possess a unique regulatory role in cyanobacteria, particularly in response to nitrogen starvation. In *Synechocystis* sp. PCC 6803 (now termed *Synechocystis*), two genes are annotated as iPGAMs, *slr1124* and *slr1945*. The product of *slr1124* was previously identified as a phosphoserine phosphatase and has an essential role in serine biosynthesis in cyanobacteria (5). Notably, we could demonstrate that the product of *slr1945* indeed is an iPGAM, and its regulation is crucial in the biosynthesis of carbon storage polymers during the adaptation to nitrogen-limiting periods (6). The acclimation response of non-diazotrophic cyanobacteria to nitrogen limitation occurs in a process termed chlorosis (7, 8). In the early phase of chlorosis, cells undergo a final doubling before cell cycle arrest, and they immediately start degrading their pigments and forming glycogen as carbon storage (9, 10). Some species also produce polyhydroxybutyrate (PHB) for an as-yet-unknown reason (11). The P_II_ signal transduction protein is a critical factor in the adaptation to nitrogen limitation. It acts as a sensor of the intracellular energy via the binding of ADP and ATP and carbon-nitrogen status via binding 2-OG. P_II_ directly or indirectly interacts with various enzymes and proteins, affecting a plethora of cellular mechanisms (12, 13). 2-OG binds to P_II_ in synergy with ATP and gives rise to a conformation of P_II_ that prevents interactions with many P_II_ target proteins (reviewed in (14)). One example of direct interaction is N-acetyl-L-glutamate kinase (NAGK), serving as a model to study P_II_-enzyme interactions (14, 15). More recently, a novel P_II_ interactor,

PirA (**P**_II_-**i**nteracting **r**egulator of **a**rginine synthesis), was discovered, whose interaction with P_II_ additionally regulates the NagK indirectly (16). Activation of the global nitrogen control transcription factor NtcA is indirectly under P_II_ control via the NtcA-co-activator PipX (**P**_II_ **i**nteracting **p**rotein **X**). Under low energy or low 2-OG conditions, P_II_ binds and sequesters PipX, whereas this complex dissociates, releasing PipX when 2-OG levels increase due to low nitrogen conditions. Then, PipX co-activates NtcA to stimulate the expression of over 80 genes required for low nitrogen acclimation(17–19). The newly discovered PirC (Sll0944, **P**_II_ **i**nteracting **r**egulator of **c**arbon metabolism) is like PipX and PirA, a small protein with no enzymatic activity. Still, it modulates the activity of a particular interaction partner, in this case, iPGAM. During nitrogen-supplemented vegetative growth, when 2-OG levels are relatively low, P_II_ forms a complex with PirC, thereby preventing iPGAM inhibition. Upon nitrogen limitation, 2-OG levels increase, and PirC dissociates from the P_II_ complex, which can then inhibit iPGAM. This blocking of the iPGAM stimulates glycogen formation and reduces carbon flow into lower glycolysis, from where many anabolic pathways, including the GS-GOGAT cycle, are derived (6).

Unlike the regulation of iPGAM in *Bacillota*, the iPGAM in cyanobacteria is regulated via the protein-protein interaction with PirC. This unique type of iPGAM regulation in cyanobacteria implies specific structural features of cyanobacterial iPGAMs. Within Cyanobacteria’s iPGAM, we identified an internal loop structure and an extended C-terminus. This study aims to clarify the role of these sub-structures in cyanobacterial iPGAM and their involvement in the PirC interaction. Therefore, we investigated the effect of these sub-structures with physiological experiments via strains containing mutated iPGAMs and biochemically with the respective purified protein variants.

## Results

### *In-silico* analysis reveals unique sub-domains in cyanobacterial iPGAM

To find out if cyanobacterial iPGAMs differ from other species, we performed a multiple sequence alignment (MSA) with a simultaneous phylogenetic analysis of 338 different iPGAMs (reviewed according to https://www.uniprot.org) using Matlab®. The tree illustrates the monophyletic evolution of cyanobacterial and red algae iPGAM, distinguishing them from all other tested iPGAMs of bacterial species, plants, and the rare metazoan iPGAMs. Intriguingly, the chloroplastic iPGAMs of red algae share a common branch with the enzymes of cyanobacteria, implying a common ancestry from the endosymbiont. In contrast, in green plants, the iPGAM of the endosymbiont was replaced. Furthermore, the tree shows a deep-rooted divergence of iPGAMs from α-cyanobacteria and β-cyanobacteria (Figure 1).

**Figure 1.**
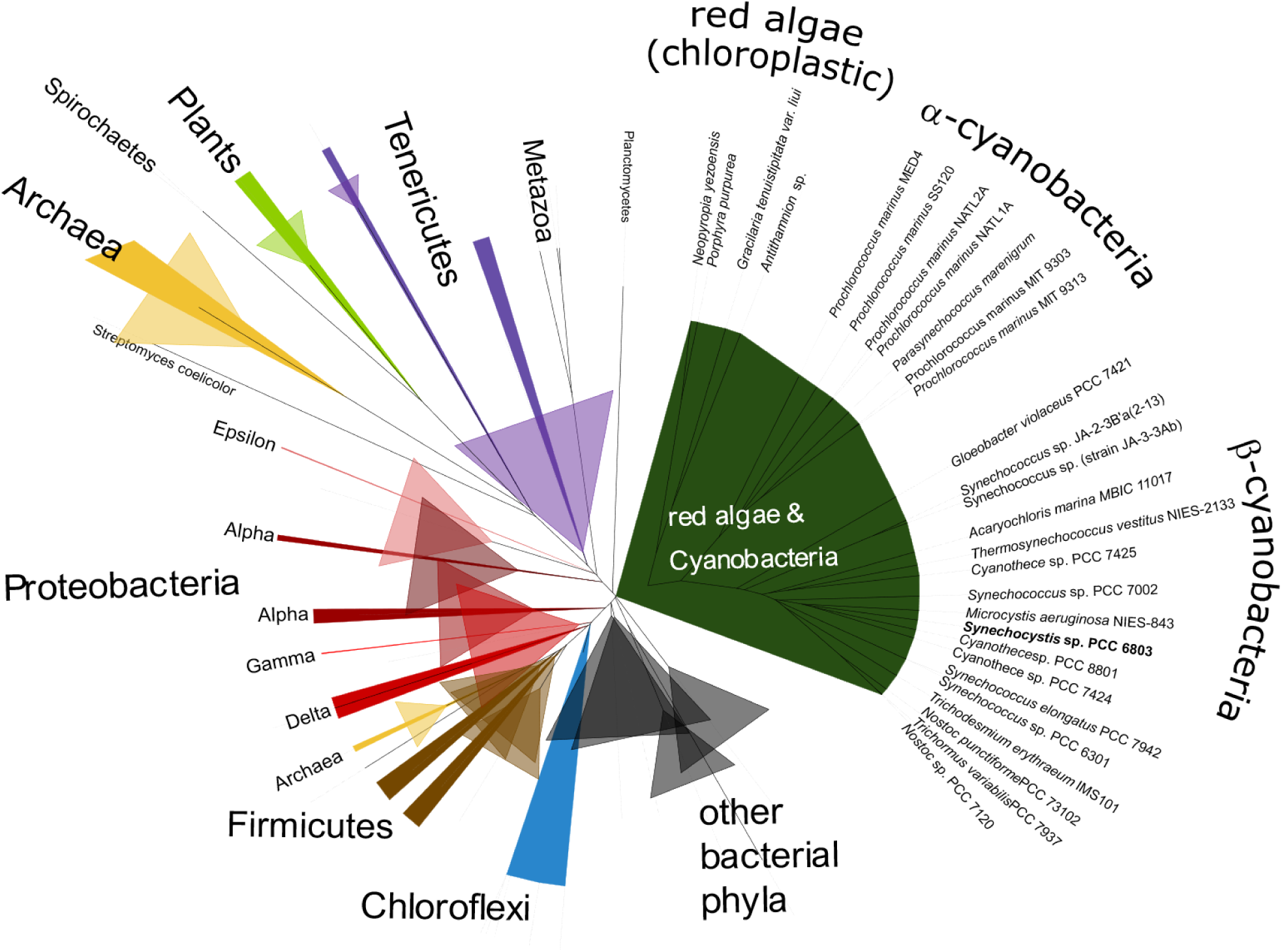
Phylogenetic tree calculated with 338 different iPGAMS (reviewed, according to UniProt).

The MSA revealed two exclusive sequence segments in cyanobacterial iPGAMs: an inner segment of 17 amino acids near the end and an extended C-Terminus (CT) (Figure S1). Accordingly, the sequence conservation of the two segments was tested based on an MSA of 644 cyanobacterial iPGAMs (Figure S 1 B & C). The inner segment comprises 17 highly conserved amino acids, which almost always start with a triplet of glutamate, glycine, and glutamate (EGE). The lysine residue at position six of the loop is also highly conserved, only replaced by arginine or glutamine and followed by hydrophobic amino acids At positions 14 and 16, there is a high probability of a hydrophobic amino acid, followed by arginine in most cases. By contrast, the CT segment, also unique to cyanobacteria, exhibits poor sequence conservation except for arginine and proline at this segment’s seventh and ninth position, respectively.

For further analysis, the *Synechocysetis* iPGAM (Slr1945) structure was predicted using SWISS-MODEL and AlphaFold 3 server to get more information on Slr1945 and the segments (Figure 2).

**Figure 2.**
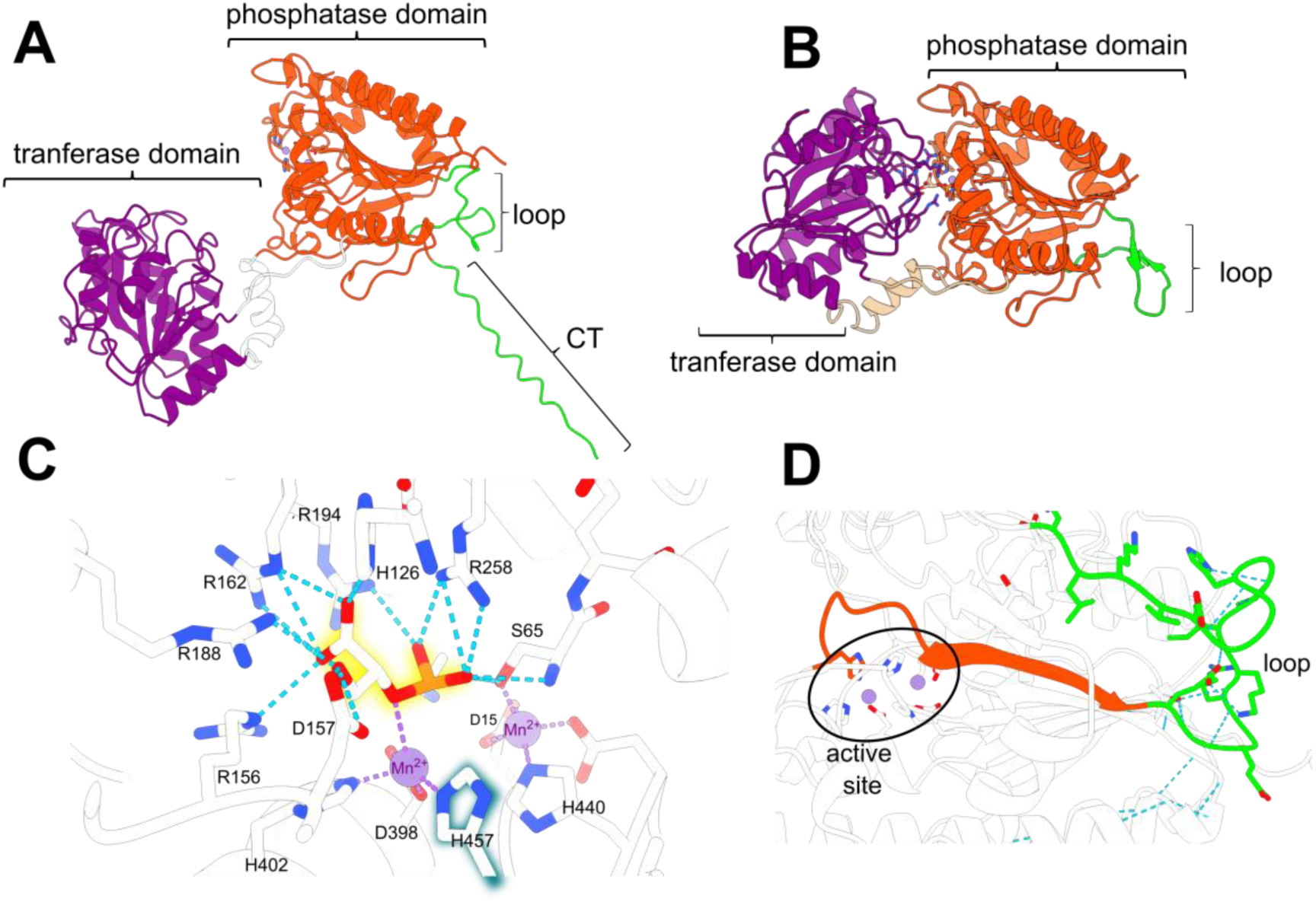
Structure of iPGAM of *Synechocystis* (Slr1945) **(A)** AlphaFold prediction of the Slr1945, prediction confidence shown in Sup. Res. Figure 2; **(B)** SWISS-MODEL prediction of the Slr1945 (PDB template: 1o98, iPGAM of *Geobacillus stearothermophilus)* Orange area = phosphatase domain, purple area = transferase domain, green areas = exclusive segments of cyanobacterial iPGAM **(C)** Contributing residues in the catalytical center of Slr1945 according to the SWISS-MODEL prediction. The numbers are based on the position in the *Synechocystis* sequence, shown in Sup. Res. Figure 1 D; **(D)** Connection of the loop to the catalytical center over the β-strand.

The structure predictions of the iPGAM of *Synechocystis* showed the typical division in the phosphatase- and transferase sub-domain of iPGAMs. They revealed a loop structure of the inner segment and a random coil at the C-terminus (CT) with local proximity (Figure 2 A & B, Figure S 2, CT only predicted by AlphaFold). Based on SWISS-MODEL prediction, the amino acid residues contributing to the active site of the *Synechocystis* iPGAM could be elucidated (Figure 2 C). The loop segment appears to be directly connected to the catalytic center, especially the histidine 457 (H457), by ten amino acids long ß-strand (Figure 2 D).

### Mass photometry indicates the involvement of three PirC monomers in the iPGAM interaction

Recombinant strep-tagged iPGAM and -PirC were produced in *E. coli* and purified by Strep-Tactin® Superflow® affinity chromatography. The purified proteins were analyzed via mass photometry to assess the oligomeric structure of iPGAM, PirC, and the iPGAM-PirC complex. The individual protein measurements used iPGAM concentrations of 10 nM. In preliminary experiments, we tested the optimal ratios for the complex measurement. The iPGAM-PirC ratio of 1:3 showed the most accurate results according to the low appearance of additional peaks (monomers of iPGAM and higher oligomers of PirC) and significant protein counts.

Mass photometry determined monomeric iPGAM particles with masses between 73-88 kDa compared to a calculated size of iPGAM of 60 kDa. For purified recombinant PirC, mass photometry determined a maximum of particles between 100 and 114 kDa, fitting to a hexameric structure of the PirC protein (strep-PirC = 14,723.35 Da). Furthermore, a shoulder of around 180 to 210 kDa indicates a higher oligomerized species of PirC. Analyzing the PirC-iPGAM complex resulted in a maximum peak of 125 kDa (replicate 2 = 134 kDa, replicate 3 = 137 kDa). Again, a shoulder appeared at the same position as in the PirC graph. With a size of 125 kDa, the peak is about 55-65 kDa larger than that of iPGAM alone, which fits the size of three PirC monomers. This suggests a complex consisting of a monomeric iPGAM to which three PirC subunits are bound. According to the results, the structure of the complex was predicted using AlphaFold (Figure 3, Figure S 4).

**Figure 3.**
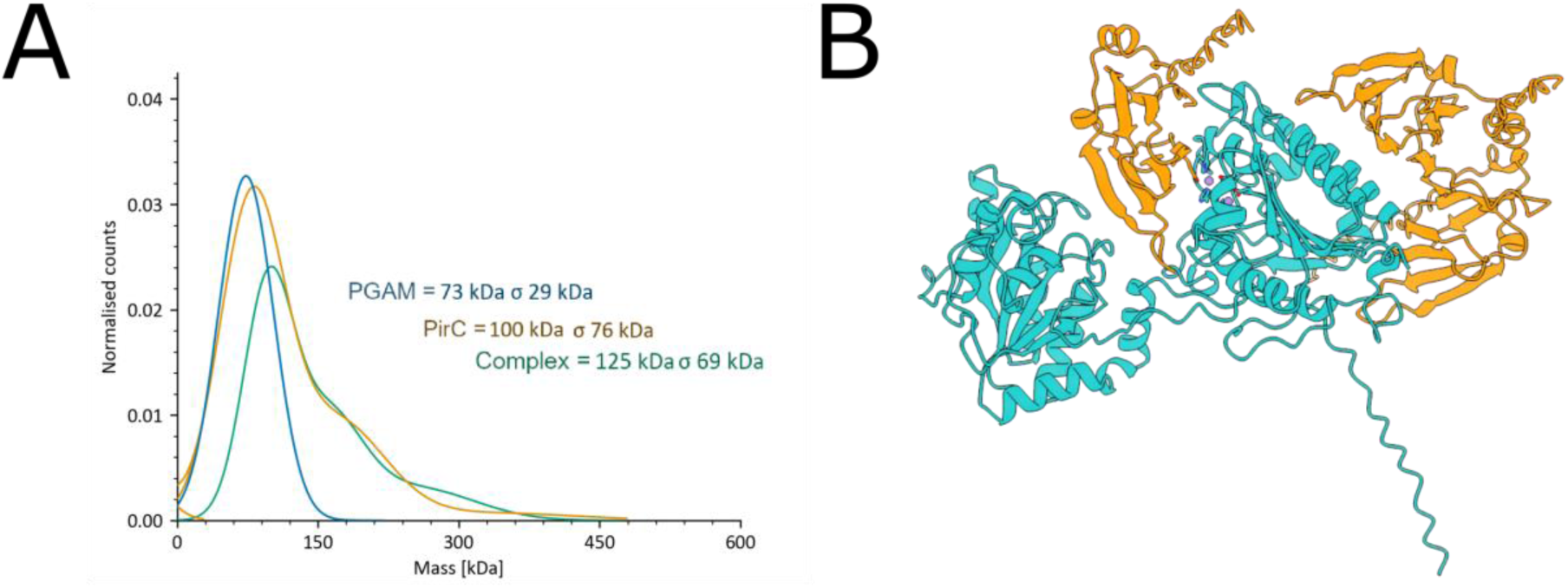
Mass photometry of strep-iPGAM, strep-PirC and their complex **(A)**, and structure of the iPGAM-PirC complex **(B)**. Representative graph of one individual measurement. Triplicates are shown in Figure S 3.

The prediction revealed three individual PirC monomers binding to iPGAM at three sites. One monomer is located in the binding cleft of the substrate between the phosphatase and transferase domain. A second monomer is located near two alpha-helices at the phosphatase domain. The third monomer also contacts the iPGAM at the phosphatase domain and is close to the loop segment. A deeper view revealed a hydrogen bond (H-bond) between highly conserved lysine 473 (K473, loop-position = 6) with a conserved glutamine residue (E468, loop-position = 1) within the loop (Figure S 5 A). In contrast, in the complex prediction of iPGAM and PirC, the tyrosine (Y39) residue of one PirC monomer also forms an H-bond with the K473, preventing the formation of the H-bond of K473-E468 (Figure S 5 B).

### Sub-structure-free variants alter the binding of PirC and inhibitory characteristics

The exclusive co-occurrence of PirC with the loop and CT segments in cyanobacteria implied functional relations. To further investigate the functional significance of these structures, three variants of the *Synechocystis* iPGAM, each with an N-terminal strep-tag, were created. First, variant iPGAM-Δloop, where the entire loop was replaced by a five amino acid sequence (TKKGI) present at homologous localization in *Geobacillus stearothermophilus* iPGAM. Second, the CT segment was deleted, creating variant iPGAM-ΔCT, and third, a combination of both alterations, iPGAM-ΔloopΔCT. In a preliminary experiment, we analyzed the interaction of the various iPGAM variants with PirC by pull-down analysis of strep-Tactin with immobilized strep-iPGAMs as bait and His-tagged PirC as analyte. Surprisingly, all three variants retained their interaction capacity with iPGAM (Figure S 6). To gain more insight into PirC binding, we performed Biolayer interferometry (BLI) measurements using an Octet® K2 system. The His-tagged version of PirC was immobilized on the surface of a Ni^2+^-NTA sensor. The different strep-iPGAM variants were then used as analytes in various concentrations for binding kinetics (Figure 4).

**Figure 4.**
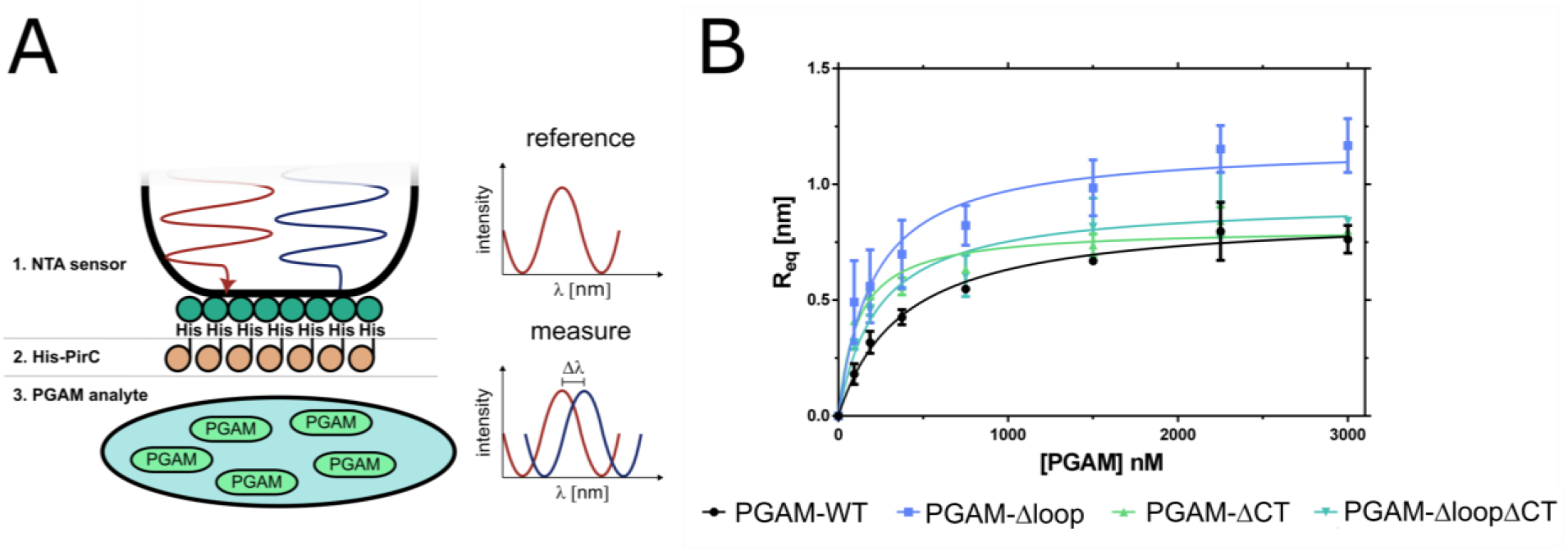
Binding of PirC and iPGAM variants **(A)**– Principle of BLI measurement with His-PirC and iPGAM variants. In BLI, binding ligands (His-PirC) and analytes (strep-PGAM) at the sensor tips alter the surface’s reflection properties, resulting in a wavelength phase shift. This shift is proportional to the bound molecule. It allows real-time detection of binding, from which the kinetic constants can be calculated **(B)** - Binding kinetics of iPGAM-WT, iPGAM-Δloop, iPGAM-ΔCT, and iPGAM-ΔloopΔCT. Each point represents a mean of technical triplicates. The error bars represent the standard deviation.

The BLI experiments revealed significant differences between the variants. Losing the loop, CT segment, or both caused an increased affinity for PirC. The K_D_ was reduced by half in the iPGAM-Δloop and iPGAM-ΔloopΔCT compared to the WT. The iPGAM-ΔCT had even a 4-times lower K_D_. The iPGAM-Δloop also had a 35 % higher binding maximum than the WT.

The BLI experiment proved that the deletion of the loop and CT structures positively affected the binding of PirC, but it did not reveal any effects on enzyme activity. Therefore, coupled enzymatic assays were carried out to check if there was a change in the catalytic properties of the iPGAM variants and the inhibitory effects exerted by PirC. First, the effect of the co-factor Mn^2+^ was tested to find the optimal manganese concentrations for each iPGAM variant (Figure S 7). The iPGAM-Δloop and the iPGAM-ΔloopΔCT required 20 times more manganese to achieve maximum activity. Therefore, a concentration of 50 µM MnCl_2_ was used in further experiments to achieve the maximum activity with all variants. First, the influence of the segment truncations on iPGAM activity was tested in the absence of PirC (Figure 5 A). Next, the experiments were repeated in the presence 50 nM, 100 nM, 200 nM, 400 nM, 800 nM, and 1600 nM PirC to gain information on the inhibition mechanism (Figure 5 C-F, only the results with or without 400 nM PirC are shown in Figure 5, Figure S 8 - S 9, kinetic parameters shown in Table S 1). In addition, the value at 0.75 mM 3-PGA for each iPGAM assay at the various PirC concentrations was used to prepare a dose-response curve to determine the IC_50_ of PirC for each iPGAM variant (Figure 5. B; Table 2). The coupling enzymes were tested with 2-PGA as a substrate to ensure that PirC does not affect the coupling reactions (Figure S 8, E). To analyze the inhibition of PirC on the different variants, the Michaelis-Menten (MM) kinetics were transformed to Hanes-Woolf kinetics (HW). The HW transformation of MM kinetics can show the strength and type of inhibition. The steeper the inhibition lines compared to the non-inhibited line, the more potent the inhibition. The convergence of the two lines without crossing before the y-axis indicates a non-competitive mode of inhibition (Figure 5). We observed this for iPGAM-WT (Figure 5. C). Additionally, the increased K_m_ of the iPGAM-WT–PirC complex indicates competitive inhibiting properties, whereas the decreased v_max_ again showed non-competitive inhibition. This implies a type of mixed inhibition by PirC (Table 2).

**Figure 5.**
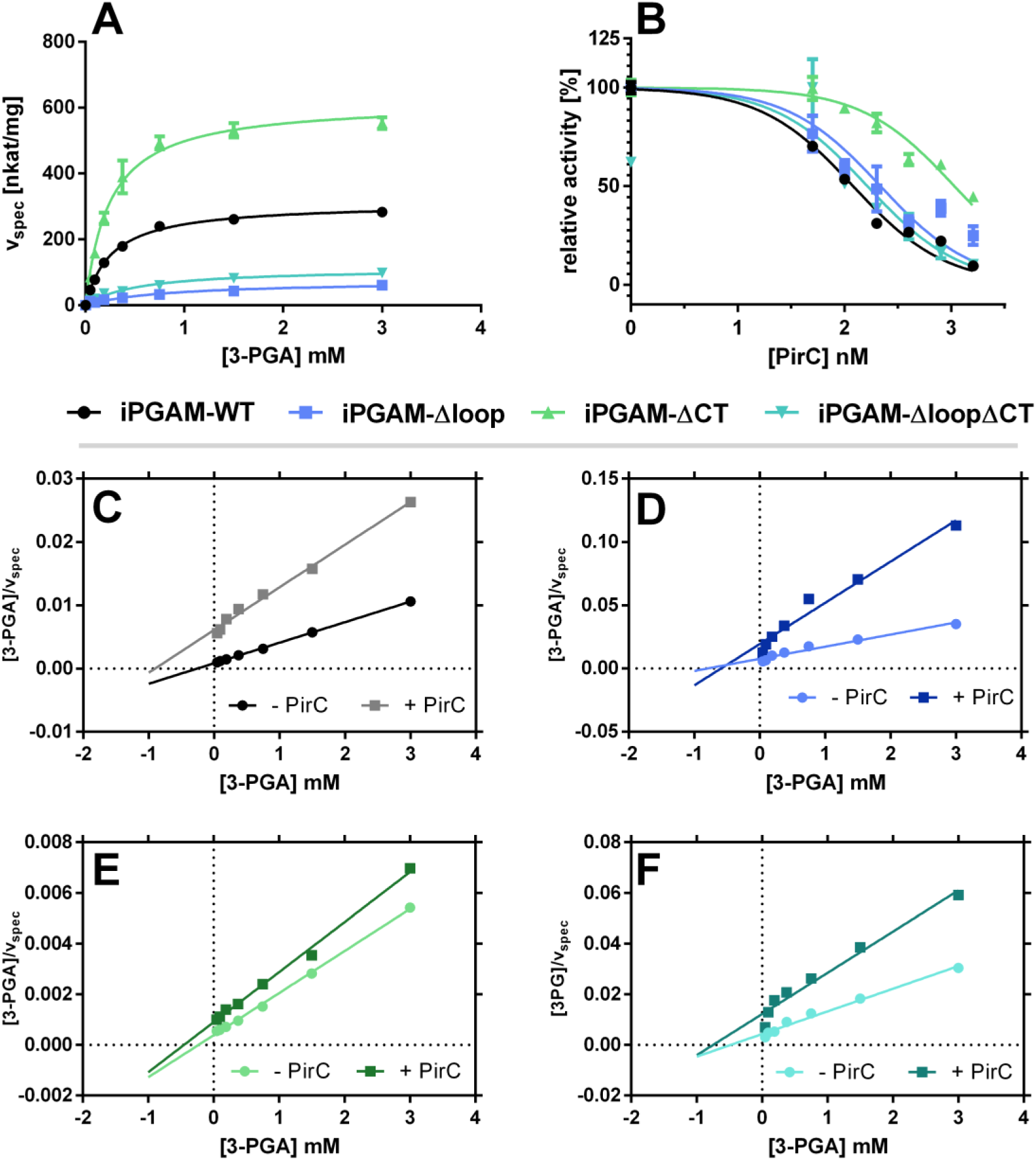
Activity of iPGAM variants of *Synechocystis* sp. PCC 6803 and the effect of PirC on activity. **(A)** Michaelis-Menten kinetics of iPGAM variants without PirC. **(B)** – Dose-response curve of PirC on the iPGAM variants at 0.75 mM 3-PGA. Each point represents the mean of three independent measured triplicates. The error bar depicts the standard deviation of the triplicate. **(C)**. - HW kinetic transformation of WT with and without PirC (400 nM) **(D)** - HW kinetic transformation of Δloop with and without PirC (400 nM). (**E)** – HW kinetic transformation of ΔCT with and without PirC (400 nM) **(F)** - Hanes-Woolf kinetic transformation of ΔloopΔCT with and without PirC (400 nM). Each point represents the transformed HW value of the mean of the triplicates. HW transformation → x-Axis = [substrate concentration], y-Axis = [substrate]/v.

**Table 1.** Binding kinetic parameters of the different iPGAM variants.

| Parameter | iPGAM-WT | iPGAM-<br>Δloop | iPGAM-<br>ΔCT | iPGAM-<br>ΔloopΔCT |
| --- | --- | --- | --- | --- |
| $R_{\max}$ [nm] | 0.872 ± 0.033 | 1.170 ± 0.063 | 0.810 ± 0.025 | 0.9291 ± 0.047 |
| $K_D$ [nM] | 378.4 ± 50.3 | 203.2 ± 45.04 | 113.6 ± 18.3 | 233.4 ± 45.3 |

**Table 2.** Kinetic parameters of iPGAM-WT, -Δloop, -ΔCT, and -ΔloopΔCT. Values represent the mean of triplicates. The error range of Km, vmax, and kcat represents the standard error calculated by GraphPad Prism. The error of kcat · Km^-1^ was calculated using error propagation—lower Part. IC50 of PirC was detected at 0.75 mM 3-PGA.

| Sample | PirC | $K_m$ [mM] | $v_{max}$ [nkat · mg<sup>-1</sup>] | $k_{cat}$ [s<sup>-1</sup>] | $k_{cat} \cdot K_m^{-1}$ [s<sup>-1</sup> · M<sup>-1</sup>] |
| --- | --- | --- | --- | --- | --- |
| iPGAM-WT | - | 0.265 ± 0.013 | 310.6 ± 4.5 | 18.6 ± 0.3 | 70,241.7 ± 3,588.3 |
| | + | 1.018 ± 0.070 | 154.4 ± 4.5 | 9.3 ± 0.3 | 9,096.3 ± 681.0 |
| iPGAM-Δloop | - | 1.087 ± 0.141 | 114.3 ± 6.4 | 6.7 ± 0.4 | 6,189.5 ± 873.9 |
| | + | 0.750 ± 0.132 | 32.1 ± 2.2 | 1.9 ± 0.1 | 2,522.3 ± 475.9 |
| iPGAM-ΔCT | - | 0.242 ± 0.019 | 617.9 ± 13.7 | 35.9 ± 0.8 | 148,468.5 ± 11,979.3 |
| | + | 0.482 ± 0.056 | 521.4 ± 20.7 | 30.3 ± 1.2 | 62,813.9 ± 7,711.8 |
| iPGAM-ΔloopΔCT | - | 0.547 ± 0.059 | 112.8 ± 4.3 | 6.4 ± 0.2 | 11,760.0 ± 1,355.4 |
| | + | 0.961 ± 0.182 | 65.7 ± 5.2 | 3.7 ± 0.3 | 3,900.7 ± 802.5 |
| Sample | | $IC_{50}$ [nM] | | | |
| iPGAM-WT | | 120.1 ± 1.06 | | | |
| iPGAM-Δloop | | 217.0 ± 1.14 | | | |
| iPGAM-ΔCT | | 1069 ± 1.07 | | | |
| iPGAM-ΔloopΔCT | | 163.5 ± 1.25 | | | |

Compared to iPGAM-WT, the iPGAM-Δloop variant had drastically reduced activity but was still inhibited by PirC. Interestingly, inhibition was less efficient, with a two-fold increase in IC_50_ for PirC compared to inhibiting the iPGAM-WT (IC_50, iPGAM-WT_ = ∼ 120 nM, IC_50, iPGAM-Δloop_ = ∼ 220 nM). More striking differences were observed for the iPGAM-ΔCT variant. First, the catalytic efficiency (CE) of iPGAM-ΔCT was severely enhanced with a CE value of 150,000 s^-1^· M^-1^ compared to ∼70,000 s^-1^· M^-1^ for WT. Furthermore, inhibition of iPGAM-ΔCT by PirC was drastically diminished, as shown by the low difference in slope between the non-inhibited and inhibited states (Figure 5. E). The inhibited iPGAM-ΔCT still had a CE of 63,000 s^-1^ · M^-1^, which is close to the activity of non-inhibited iPGAM-WT. The dose-response curves of PirC and the calculated IC_50_ demonstrated a 10-fold decreased efficiency of PirC to inhibit iPGAM-ΔCT compared to iPGAM-WT (IC_50, iPGAM-WT_ = ∼ 120 nM, IC_50, iPGAM-ΔCT_= ∼ 1070 nM). The iP-GAM-loopΔCT variant combined properties of the two single mutations: Deleting the C-terminus in the iPGAM-Δloop variant increased its catalytic efficiency and partially decreased the inhibitory effect of PirC compared to the Δloop variant (Table 2).

### Sub-domain-free variants influence the physiology during chlorosis

To investigate the effect of the above-described alterations of the iPGAM variants on the physiology of *Synechocystis*, mutant strains were generated by homologous recombination of native iPGAM with the iPGAM variants followed by spectinomycin (Δloop) or chloramphenicol (ΔCT and ΔloopΔCT) as selection markers. As controls, the WT was used as a reference for the standard interaction between iPGAM and PirC and a PirC-deletion mutant (ΔPirC) for native iPGAM without PirC regulation.

Since the PirC-iPGAM interaction affects physiology during nitrogen limitation, we tested the effect in nitrogen deprivation experiments, in which we analyzed the OD_750_, the glycogen amount, and the PHB levels after 14 days. As control experiments, we also tested the effect of nitrogen-rich conditions with nitrate or ammonia (Figure 6).

**Figure 6.**
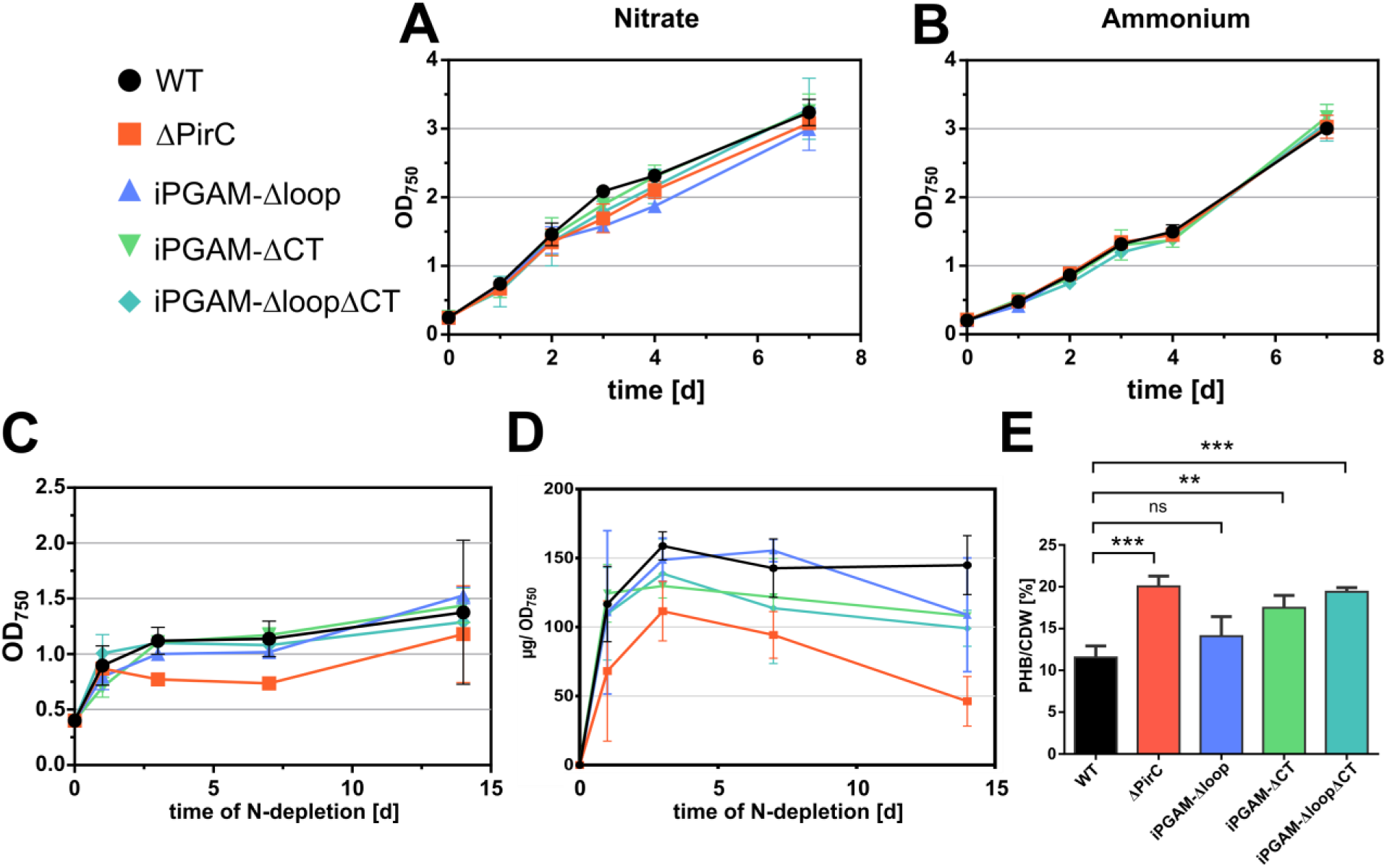
Effect of iPGAM-Δloop, -ΔCT, and -ΔloopΔCT on the physiology of *Synechocystis* sp. PCC 6803. (**A)** Growth curves of iPGAM variants with 17 mM NaNO4 (**B)** Growth curves of iPGAM variants with 5 mM NH4Cl. (**C)** OD750 of iPGAM variants during nitrogen depletion. (**D)**. Glycogen content in iPGAM variant strains during nitrogen depletion. Each point represents the mean, and the error bars represent the standard deviation of independent triplicates. (**E)** PHB content after 14 days of nitrogen depletion in the iPGAM variants. Each bar and error bar represents the mean of biological triplicates.

All tested were unaffected in nitrogen-supplemented vegetative growth. How-ever, the ΔPirC strain could not carry out the typical final cell division upon nitrogen starvation, unlike the iPGAM variants and WT cells. The ΔPirC accumulated less glycogen and significantly increased levels of PHB compared to the WT. The iPGAM-Δloop mutant accumulated glycogen at similar levels as the WT but required more time to reach its maximum level. The iPGAM-ΔCT and -ΔloopΔCT strains showed intermediary phenotypes between ΔPirC and WT for glycogen accumulation. A hallmark of the *pirC* mutation was the enormously increased accumulation of PHB. In this respect, the iPGAM-ΔCT and - ΔloopΔCT strains resembled the ΔPirC. After 14 days of chlorosis, the PHB level of iPGAM-ΔCT (18.2 ± 0.4 mg · CDW^-1^) and ΔloopΔCT (19.4 ± 0.3 mg · CDW^-1^) strains reached PHB levels similar to the high producer ΔPirC (20.1 ± 1.0 mg · CDW^-1^). Meanwhile, the iPGAM-Δloop strain (12.9 ± 0.5 mg · CDW^-1^) only reached PHB amounts, which were insignificantly higher than levels reached within the WT (11.5 ± 1.5 mg · CDW^-1^).

## Discussion

This research unveils a unique insight into the pivotal role of the 2,3-bisphos-phoglycerate-independent phosphoglycerate mutase (iPGAM) in regulating carbon flux in cyanobacteria. The key player, PirC, inhibits the iPGAM reaction upon nitrogen starvation, redirecting the carbon flux from lower glycolysis to glycogen formation. Release from a tight complex with P_II_ upon perceiving nitrogen starvation initiates the formation of the PirC-PGAM complex. (6). The small regulatory protein PirC is a distinctive feature of cyanobacteria. Our bioinformatics analysis revealed two structural elements exclusively in cyanobacterial iPGAM, indicating a unique functional connection with PirC. The phylogenetic analysis showed that cyanobacterial iPGAMs diverged early from iPGAMs of other species. Notably, the iPGAMs of red algae show a phylogenetic relation to cyanobacterial iPGAM despite lacking the two characteristic segments. The chloroplasts of red algae have retained several features from the early endosymbiont, which were later lost in the evolution of green plants, such as components of the light-harvesting system. Furthermore, red algae exhibit P_II_ signaling features that resemble cyanobacteria, such as N-Acetyl-L-Glutamate kinase regulation by P_II_ (20). The phylogenetic tree of cyanobacterial iPGAM corresponds strikingly well with a recent whole genome phylogeny of cyanobacteria (21), which considers many more genes than the usually used 120 housekeeping genes (22). If this refined phylogenetic tree accurately reflects the evolution of cyanobacteria, it suggests that the characteristic features of iPGAM and their regulation by PirC emerged early in cyanobacterial evolution. Alternatively, the prominent divergence of iPGAM homology between α- and β-cyanobacteria would indicate a different trend in protein evolution in these cyanobacterial groups. In red algae, the evolutionary loss of PirC led to the disappearance of the CT and loop segments, but other features of iPGAM were conserved, maintaining the phylogenetic clustering. In contrast, the evolution towards green plants involved significantly streamlining the P_II_ signaling system along with metabolic rearrangements. The endosymbiont’s iPGAM was replaced by an unrelated enzyme, not associated with P_II_ signaling, along with the translocation of the *glnB* gene to the nucleus and the reorganization of the P_II_ signaling system (such as the acquisition of the glutaminesensing C-terminal domain).

Detailed kinetic analysis of iPGAM now shows characteristics of both competitive- and non-competitive inhibition by PirC. The enzyme has a lowered affinity to the substrate in the presence of PirC, as demonstrated by a lowered K_m_. Furthermore, the iPGAM also has a reduced maximal activity in the presence of PirC (shown by v_max_ and k_cat_). Roychowdhury *et al.,* 2015 showed that iP-GAMs have an open conformation when no substrate is bonded (as seen also in our AlphaFold prediction). The substrate diffuses into the binding pocket and induces the closing of this cleft by phosphatase- and transferase domain movement, which is required for the reaction to occur (23). The PirC peptide, which AlphaFold predicted to be localized in the phosphatase-transferase interdomain cleft, possibly hinders the cleft closure and, thereby, the catalytic reaction.

The exchange of the loop drastically reduced the activity of the iPGAM. The iPGAM-Δloop only reached one-third of the maximal activity as the iPGAM-WT, and the changed K_m_ evidences a four times loss in substrate affinity. The H457, connected to the loop, is equivalent to H462 from the *G. stearothermophilus* iPGAM, which contributes to the binding of manganese (Figure 2 C) (24). In agreement, the iPGAM-Δloop variant, where H457 is deleted, has a reduced affinity to Mn^2+^ (K_half, WT_(Mn^2+^) = ∼1.3 µM; K_half, Δloop_(Mn^2+^) = ∼10.5 µM), and the enzyme requires much higher Mn^2+^ concentrations to reach its maximal activity. The AlphaFold prediction of the iPGAM structure revealed a H-bond between K473 and E468 within the loop. This interaction possibly stabilizes the β-strand connection to the catalytic center, favoring the manganese binding. The iPGAM of *G. stearothermophilus,* which does not possess this loop, requires manganese concentrations that are 1,000-fold higher than the K_half_(Mn^2+^) of *Synechocystis* iPGAM for maximal activity (25). The low activity of the PGAM-Δloop indicates that the lowered manganese affinity consequently affects the whole reaction. Hence, the H457 residue contributes to the coordination of the bond Mn^2+^, which is essential in the phosphatase reaction of the enzyme. The H-bond K473-E468 within the loop structure keeps the H457 in an optimal position, and Mn^2+^ binds with higher affinity. Furthermore, the Mn^2+^ coordinates the phosphoester bond of 2- or 3-PGA and thus enables the hydrolyzation of the phosphate. In the iPGAM-Δloop variant, the K473-E468 interaction does not exist, and H457 could be in another orientation within the catalytic center, resulting in a lowered K_m_. PirC also prevents the K473-E468 interaction. One of three binding sites of PirC is located toward the surface of the phosphatase sub-domain in direct connection to the loop of the iPGAM. Here, the Y39 of PirC is predicted to form a hydrogen bond with K473, which consequently prevents the K473-E468 H-bond. This also changes the H457 orientation and leads to the inhibition of the iPGAM. Intriguingly, the iPGAM-Δloop variant has a higher affinity for PirC, indicating a conformational change imposed by the loop deletion that facilitates PirC interaction, confirming the structural prediction that the loop segment is not a prerequisite for PirC binding.

In contrast to the iPGAM-Δloop, the iPGAM-ΔCT has a two-fold increased v_max_ compared to iPGAM-WT, whereas the K_m_ is not altered. At the same time, the affinity towards PirC is strongly enhanced, but PirC only weakly inhibits the iPGAM reaction: The inhibitory effect of PirC on iPGAM-ΔCT activity is 10-fold lower than on iPGAM-WT (IC_50, PirC_(iPGAM-WT) = ∼ 120 nM, IC_50, PirC_(iPGAM-ΔCT) = ∼ 1070 nM). This directly implies that the C-terminal flexible extension plays a crucial role in modulating the activity of iPGAM and transmits the inhibitory effect of PirC binding to the catalytic center. Possibly, this C-terminal extension lowers the v_max_ of the reaction by interfering with the domain closure. By removing this tail, the enzyme can work at its maximum pace. In agreement, the binding of PirC would place the C-terminal tail in a position where the inhibitory effect is augmented. Without the C-terminal extension, the binding of PirC cannot exert this inhibitory function but facilitates its binding to the iPGAM body. This can be explained by the assumption that the binding of the CT extension by PirC is thermodynamically unfavorable. Residual inhibition of the iPGAM-ΔCT variant requires 10-fold higher concentrations of PirC, although the affinity of these partners has increased, which appears counterintuitive. It suggests that the binding of PirC protomers to low-affinity binding sites of iPGAM-ΔCT is required to achieve inhibition. These additional binding events may impair the catalytically driven domain closure of iPGAM. Concerning the iPGAM-Δloop-ΔCT variant, it shows a similar increased K_m_ for Mn^2+^ as the single iPGAM-Δloop, which agrees with the apparent role of the loop segment in high Mn^2+^ affinity. Surprisingly, however, the combined removal of both the loop and the CT segment gave rise to compensatory effects concerning its catalytic properties, as the catalytic efficiency is in between that of iPGAM-Δloop and iPGAM-WT variants and inhibition of enzyme activity by PirC affects the K_m_ in a similar way as deletion of the CT segment. Overall, the binding of PirC to different sites of iPGAM, sites near the unique loop structure, and the CT segment is likely the structural basis of the observed mixed-type inhibition. Structural analysis is necessary to resolve further molecular details of PirC-mediated inhibition.

Under nitrogen-replete conditions, all the iPGAM variant strains grow similarly to the WT. This shows that during nutrient-replete vegetative growth, the low activities of the iPGAM-Δloop and iPGAM-ΔloopΔCT variants are sufficient to maintain metabolic homeostasis. Conversely, the excessive activity of the iP-GAM-ΔCT variant during vegetative growth has neither a positive nor a negative effect on *Synechocystis* growth. However, a distinct phenotype of iPGAM variant strains could be observed during nitrogen starvation. Unlike the ΔPirC strain, the iPGAM-Δloop, -ΔCT, and -ΔloopΔCT strains carry out a final cell division upon shifting to a nitrogen-depleted medium. Furthermore, all iPGAM strains formed similar amounts of glycogen, whereas the ΔPirC strain is strongly affected in glycogen accumulation, in agreement with earlier observations (6). During prolonged chlorosis, the iPGAM-Δloop, -ΔCT, and -ΔloopΔCT strains showed a slightly increased glycogen degradation compared to the WT. Previously, we showed that the levels of 3-PGA, a key activator of glycogen formation, in both the WT and ΔPirC doubled within the first six hours after N-depletion (6). This indicates that the initial 3-PGA accumulation is independent of the iPGAM-PirC interaction, which also explains the initial glycogen increase in ΔPirC. During further nitrogen starvation, expression of PirC is strongly induced (26). This leads to a pronounced inhibition of iPGAM and, consequently, a further increase of 3-PGA levels, which is no longer observed in the ΔPirC. Consequently, glycogen levels continue to increase in the WT to their maximum levels, whereas it slows down in ΔPirC.

During prolonged nitrogen starvation, glycogen is slowly converted into PHB (27). In the ΔPirC strain, lack of iPGAM inhibition during chlorosis leads to accelerated carbon flow into lower glycolysis, finally resulting in PHB formation. Both strains expressing the CT-truncated iPGAM (iPGAM-ΔCT and iP-GAM-ΔloopΔCT) show almost the same amount of PHB accumulation after 14 days of nitrogen starvation as the ΔPirC. This can very likely be attributed to the reduced inhibition of these variants by PirC. In contrast, the iPGAM-Δloop strain shows similar levels to the WT, which agrees with the low activity of the iPGAM-Δloop-PirC complex.

Although the iPGAM-ΔCT expressing strains produce similar amounts of PHB during prolonged chlorosis as the PirC deficient strain. They are still able to respond to nitrogen starvation in a similar way to the WT by accumulating glycogen and performing a final cell division, whereas, in the absence of PirC, the cells go immediately into growth arrest. This could indicate additional roles of PirC beyond iPGAM inhibition for the acclimation towards nitrogen starvation. However, concerning the effect on PHB accumulation, inhibition of iPGAM seems to be the most important function. Previously, Koch *et al.* 2020 (28) achieved higher PHB levels with the ΔPirC strain by introducing additional *phaA* (acetyl-CoA acetyltransferase) and *phaB* (acetoacetyl-CoA reductase) genes, whose products catalyze the initial steps of PHB synthesis. A similar approach should also increase the PHB amounts in iPGAM-ΔCT or the iPGAM-ΔloopΔCT. In contrast to ΔPirC, these strains do not have the disadvantage of biomass loss and poorer viability during nitrogen starvation, which seems beneficial for biotechnological applications.

## Material & Methods

Detailed descriptions of the methods are shown in Supplemental Material & Methods.

### Multiple alignments and phylogenetic tree calculation of phosphoglycerate mutases

The alignments were done with Matlab®. The resulting tree was visualized with *iTol*(29).

### Structure predictions

The structure of the iPGAM was predicted using the SWISS-Model workspace and AlphaFoldServer (30–32). AlphaFold was also used to predict the structure of the complex.

### Molecular Cloning and mutagenesis

Gibson Assembly (GA) and mutagenesis PCR were used to create the plasmids. The GA was done according to the manufacturer protocol (NEB E2611S/L, E5510S).

According to the manufacturer protocol, iPGAM was mutated with the Q5® Site-Directed Mutagenesis Kit (NEB, E0554).

### Plasmid and strains

Plasmids and strains created and used in this study are listed in Table S1 and Table S 2.

### Cultivation of Cyanobacteria

Growth experiments and precultures of *Synechocystis* were cultivated in BG11, and the composition was explained by M. Mager *et al.* (33). Standard cultivation was performed at 28 °C with continuous shaking at 125 rpm at constant illumination (24 h/d, ∼50 μE m^−2^ s^−1^). The BG11 was adjusted for different experiments, as explained in the supplemental material.

For nitrogen deficiency experiments, pre-cultures of *Synechocystis* at an OD750 of 0.6-1 were washed with and resuspended in BG_11,_0 medium, and the nitrogen-free culture was inoculated to OD750 of 0.4.

Escherichia coli cultures were grown on LB medium and agar.

### Expression and Purification of Proteins

*E. coli* Lemo21(DE3) was used to overexpress proteins induced depending on the vector either by 400 mM IPTG or 200 µg anhydrotetracycline. The histagged proteins were purified using HisTrap HP columns (Cytiva, Marlborough, USA) and strep-tagged protein using the Strep-tactin® superflow columns (IBA Lifescience, Göttingen, Germany) by affinity chromatography.

### Mass photometry using the Refyn OneMP

A mass photometry experiment was used to study the variants’ oligomerization and the stoichiometry of the iPGAM-PirC complex. For this purpose, the Refyn OneMP was used. The data was analyzed using the DiscoverMP® software.

### BLI using the Octet K2 System

*In vitro* binding studies were done using bio-layer interferometry (BLI) using the Octet K2 system (Sartorius, Göttingen, Germany) according to the Bioprotocol (34).

### Phosphoglycerate Mutase Assay

The iPGAM activity was determined by a coupled enzyme assay as adapted as described previously (6).

### Glycogen Measurement

The glycogen content was quantified according to previous studies from 2 ml *Synechocystis* culture samples (35).

### PHB Quantification

Polyhydroxybutyrate (PHB) was detected using high-performance liquid chromatography (HPLC) as described previously (6, 11, 28).

## Author Contribution

Conceptualization, TO and KF; Methodology and interpretation of results, TO and KF; Overall Investigations, TO; BLI Measurements, TO & NB; Construction of pJA01 and construction of ΔPirC Strain, JA; PHB analysis, PF; Writing-original draft preparation, TO and KF; Writing-review & editing, TO, JA, PF, NB, and KF; Supervision, KF; All authors read and approved the final manuscript.

## Acknowledgments

We thank the Weir Lab at Friedrich-Miescher Laboratory at the Max-Planck Institute in Tübingen for allowing us to use their ReFeyn OneMP mass photometer. We want to thank Sofía Doello for the introduction to the instrument. The work was supported by a grant from the Deutsche Forschungsgemeinschaft (DFG) Fo195/21-1 and infrastructural support through the Cluster of Excellence EXC 2124 (Controlling Microbes to Fight Infections, CMFI, grant 390838134) at the Eberhard Karls Universität Tübingen

## References

1. Galperin MY, Bairoch A, Koonin E V. 1998. A superfamily of metalloenzymes unifies phosphopentomutase and cofactor-independent phosphoglycerate mutase with alkaline phosphatases and sulfatases. Protein Sci 7:1829.

2. Jedrzejas MJ. 2000. Structure, function, and evolution of phosphoglycerate mutases: comparison with fructose-2,6-bisphosphatase, acid phosphatase, and alkaline phosphatase. Prog Biophys Mol Biol 73:263–287.

3. Kuhn NJ, Setlow B, Setlow P. 1993. Manganese(II) Activation of 3-Phosphoglycerate Mutase of Bacillus megaterium: pH-Sensitive Interconversion of Active and Inactive Forms. Arch Biochem Biophys 306:342–349.

4. Loshon CA, Setlow P. 1993. Levels of small molecules in dormant spores of *Sporosarcina* species and comparison with levels in spores of *Bacillus* and *Clostridium* species. Can J Microbiol 39:259–262.

5. Klemke F, Baier A, Knoop H, Kern R, Jablonsky J, Beyer G, Volkmer T, Steuer R, Lockau W, Hagemann M. 2015. Identification of the light-independent phosphoserine pathway as an additional source of serine in the cyanobacterium synechocystis sp. PCC 6803. Microbiology (United Kingdom) 161:1050–1060.

6. Orthwein T, Scholl J, Spät P, Lucius S, Koch M, Macek B, Hagemann M, Forchhammer K. 2021. The novel PII-interactor PirC identifies phosphoglycerate mutase as key control point of carbon storage metabolism in cyanobacteria. Proc Natl Acad Sci U S A 118:e2019988118.

7. Allen MM, Smith AJ. 1969. Nitrogen chlorosis in blue-green algae. Arch Mikrobiol 69:114–120.

8. Neumann N, Doello S, Forchhammer K. 2021. Recovery of Unicellular Cyanobacteria from Nitrogen Chlorosis: A Model for Resuscitation of Dormant Bacteria. Microb Physiol 31:78–87.

9. Sauer J, Schreiber U, Schmid R, Völker U, Forchhammer K. 2001. Nitrogen Starvation-Induced Chlorosis in Synechococcus PCC 7942. Low-Level Photosynthesis As a Mechanism of Long-Term Survival. Plant Physiol 126:233.

10. Schlebusch M, Forchhammer K. 2010. Requirement of the Nitrogen Starvation-Induced Protein Sll0783 for Polyhydroxybutyrate Accumulation in Synechocystis sp. Strain PCC 6803. Appl Environ Microbiol 76:6101.

11. Koch M, Berendzen KW, Forchhammer K. 2020. On the Role and Production of Polyhydroxybutyrate (PHB) in the Cyanobacterium Synechocystis sp. PCC 6803. Life 10.

12. Forchhammer K, Schwarz R. 2019. Nitrogen chlorosis in unicellular cyanobacteria – a developmental program for surviving nitrogen deprivation. Environ Microbiol 21:1173–1184.

13. Forchhammer K, Selim KA. 2020. Carbon/nitrogen homeostasis control in cyanobacteria. FEMS Microbiol Rev 44:33–53.

14. Forchhammer K, Lüddecke J. 2016. Sensory properties of the PII signalling protein family. FEBS J 283:425–437.

15. Rozbeh R, Forchhammer K. 2021. Split NanoLuc technology allows quantitation of interactions between PII protein and its receptors with unprecedented sensitivity and reveals transient interactions. Scientific Reports 2021 11:1 11:1–13.

16. Bolay P, Rozbeh R, Isabel Muro-Pastor M, Timm S, Hagemann M, Florencio FJ, Forchhammer K, Klähn S. 2021. The novel pii-interacting protein pira controls flux into the cyanobacterial ornithine-ammonia cycle. mBio 12.

17. Giner-Lamia J, Robles-Rengel R, Hernández-Prieto MA, Isabel Muro-Pastor M, Florencio FJ, Futschik ME. 2017. Identification of the direct regulon of NtcA during early acclimation to nitrogen starvation in the cyanobacterium Synechocystis sp. PCC 6803. Nucleic Acids Res 45:11800–11820.

18. Forcada-Nadal A, Bibak S, Salinas P, Contreras A, Rubio V, Llácer JL. 2024. Structural understanding of NtcA regulation and of its coactivation by the adaptor PII/NtcA shuttling protein PipX, which connects PII regulation with gene expression regulation. bioRxiv 2024.08.29.607757.

19. Forcada-Nadal A, Llácer JL, Contreras A, Marco-Marín C, Rubio V. 2018. The PII-NAGK-PipX-NtcA regulatory axis of cyanobacteria: A tale of changing partners, allosteric effectors and non-covalent interactions. Front Mol Biosci 5:421097.

20. Selim KA, Ermilova E, Forchhammer K. 2020. From cyanobacteria to Archaeplastida: new evolutionary insights into PII signalling in the plant kingdom. New Phytol 227:722–731.

21. Strunecký O, Wachtlová M, Koblížek M. 2021. Whole genome phylogeny of Cyanobacteria documents a distinct evolutionary trajectory of marine picocyanobacteria. bioRxiv 2021.05.26.445609.

22. Strunecký O, Ivanova AP, Mareš J. 2023. An updated classification of cyanobacterial orders and families based on phylogenomic and polyphasic analysis. J Phycol 59:12–51.

23. Roychowdhury A, Kundu A, Bose M, Gujar A, Mukherjee S, Das AK. 2015. Complete catalytic cycle of cofactor-independent phosphoglycerate mutase involves a spring-loaded mechanism. FEBS J 282:1097–1110.

24. Jedrzejas MJ, Chander M, Setlow P, Krishnasamy G. 2000. Structure and mechanism of action of a novel phosphoglycerate mutase from Bacillus stearothermophilus. EMBO J 19:1419.

25. Chander M, Setlow P, Lamani E, Jedrzejas MJ. 1999. Structural studies on a 2,3-diphosphoglycerate independent phosphoglycerate mutase from Bacillus stearothermophilus. J Struct Biol 126:156–165.

26. Muro-Pastor MI, Cutillas-Farray Á, Pérez-Rodríguez L, Pérez-Saavedra J, Vega-De Armas A, Paredes A, Robles-Rengel R, Florencio FJ. 2020. CfrA, a Novel Carbon Flow Regulator, Adapts Carbon Metabolism to Nitrogen Deficiency in Cyanobacteria. Plant Physiol 184:1792–1810.

27. Koch M, Doello S, Gutekunst K, Forchhammer K. 2019. PHB is Produced from Glycogen Turn-over during Nitrogen Starvation in Synechocystis sp. PCC 6803. Int J Mol Sci 20:1942.

28. Koch M, Bruckmoser J, Scholl J, Hauf W, Rieger B, Forchhammer K. 2020. Maximizing PHB content in Synechocystis sp. PCC 6803: a new metabolic engineering strategy based on the regulator PirC. Microb Cell Fact 19:1–12.

29. Letunic I, Bork P. 2024. Interactive Tree of Life (iTOL) v6: recent updates to the phylogenetic tree display and annotation tool. Nucleic Acids Res 52:W78–W82.

30. Guex N, Peitsch MC, Schwede T. 2009. Automated comparative protein structure modeling with SWISS-MODEL and Swiss-PdbViewer: A historical perspective. Electrophoresis 30:S162–S173.

31. Waterhouse A, Bertoni M, Bienert S, Studer G, Tauriello G, Gumienny R, Heer FT, De Beer TAP, Rempfer C, Bordoli L, Lepore R, Schwede T. 2018. SWISS-MODEL: homology modelling of protein structures and complexes. Nucleic Acids Res 46:W296.

32. Abramson J, Adler J, Dunger J, Evans R, Green T, Pritzel A, Ronneberger O, Willmore L, Ballard AJ, Bambrick J, Bodenstein SW, Evans DA, Hung CC, O’Neill M, Reiman D, Tunyasuvunakool K, Wu Z, Žemgulytė A, Arvaniti E, Beattie C, Bertolli O, Bridgland A, Cherepanov A, Congreve M, Cowen-Rivers AI, Cowie A, Figurnov M, Fuchs FB, Gladman H, Jain R, Khan YA, Low CMR, Perlin K, Potapenko A, Savy P, Singh S, Stecula A, Thillaisundaram A, Tong C, Yakneen S, Zhong ED, Zielinski M, Žídek A, Bapst V, Kohli P, Jaderberg M, Hassabis D, Jumper JM. 2024. Accurate structure prediction of biomolecular interactions with AlphaFold 3. Nature 2024 630:8016 630:493–500.

33. Mager M, Pineda Hernandez H, Brandenburg F, López-Maury L, McCormick AJ, Nürnberg DJ, Orthwein T, Russo DA, Victoria AJ, Wang X, Zedler JAZ, Dos Santos FB, Schmelling NM. 2023. Interlaboratory Reproducibility in Growth and Reporter Expression in the Cyanobacterium Synechocystis sp. PCC 6803. ACS Synth Biol 12:1823–1835.

34. Orthwein T, Huergo LF, Forchhammer K, Selim KA. 2021. Kinetic analysis of a protein-protein complex to determine its dissociation constant (kd) and the effective concentration (ec50) of an interplaying effector molecule using bio-layer interferometry. Bio Protoc 11.

35. Doello S, Klotz A, Makowka A, Gutekunst K, Forchhammer K. 2018. A Specific Glycogen Mobilization Strategy Enables Rapid Awakening of Dormant Cyanobacteria from Chlorosis. Plant Physiol 177:594–603.

