## Supplemental Material for "Structural elements of cyanobacterial co-factor-independent phosphoglycerate mutase that mediate regulation by PirC"

### Supplementary Results

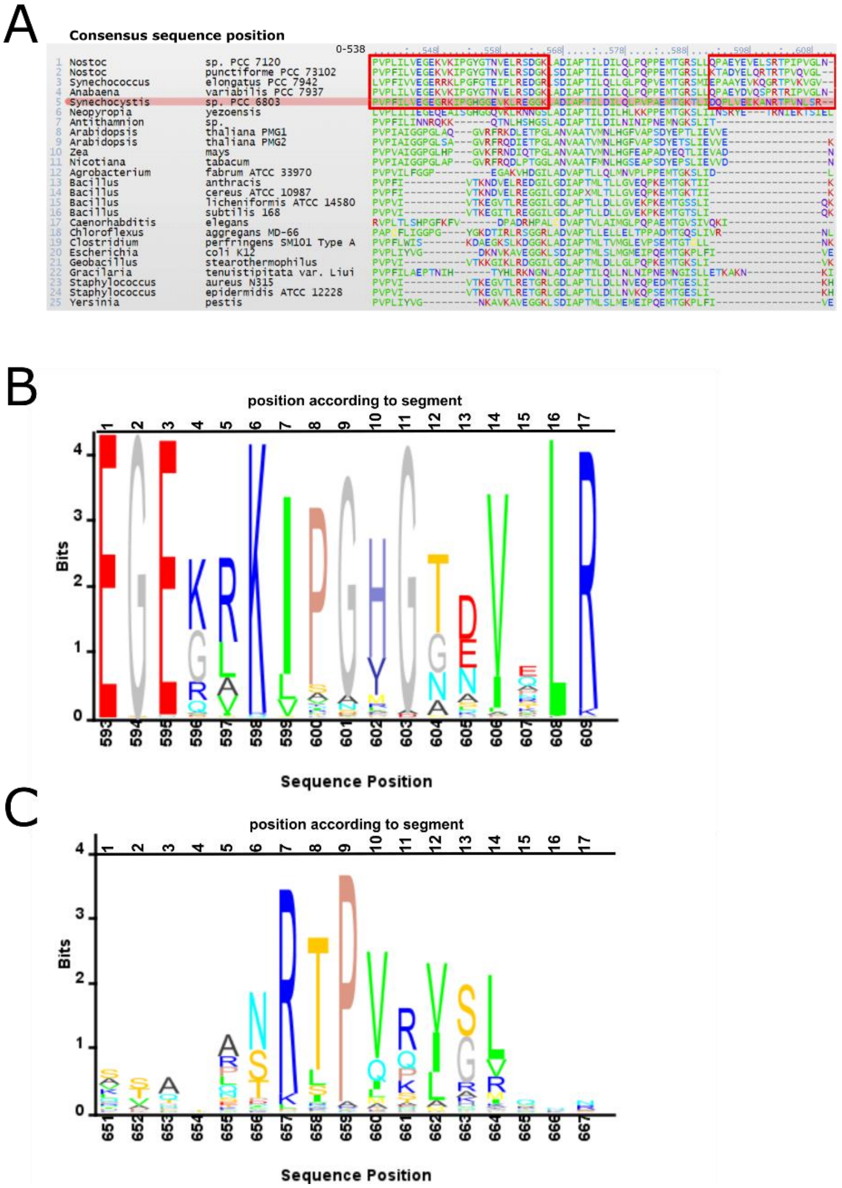


Figure S 1 – Comparison of 338 reviewed iPGAM sequences (uniprot.org) and 644 iPGAM sequences of cyanobacteria revered as active (uniprot.org). (A) Excerpt of an alignment of 338 different iPGAMs. Cyanobacteria have two unique sequences, which are highlighted by a red box. The first is called a loop according to the predicted structure. Second, an extended C-terminus. (B) The sequence logo of the loop sequence was calculated via an alignment of iPGAMs of 644 different cyanobacteria. (C) Sequence logo of the extended C-terminus in cyanobacteria.

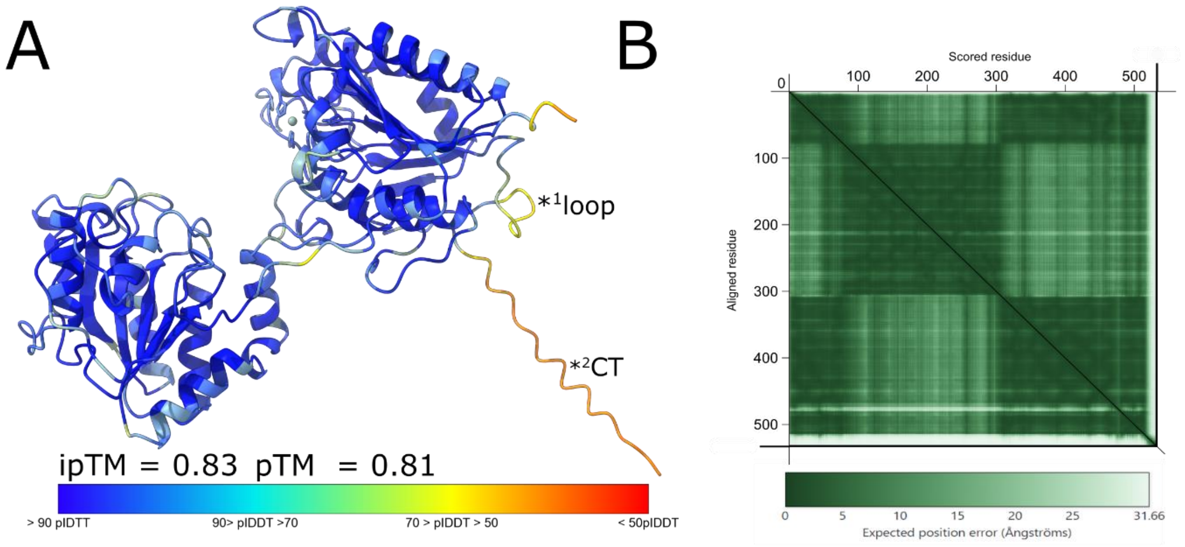


Figure S 2 – Structure of a *Synechocystis i*PGAM (A) AlphaFold-predicted structure of the PGAM colorized by the predicted IDDT matching score (pIDDT) of the structure (B) Predicted aligned error (PAE) diagram of the PGAM-PirC(tri) complex.

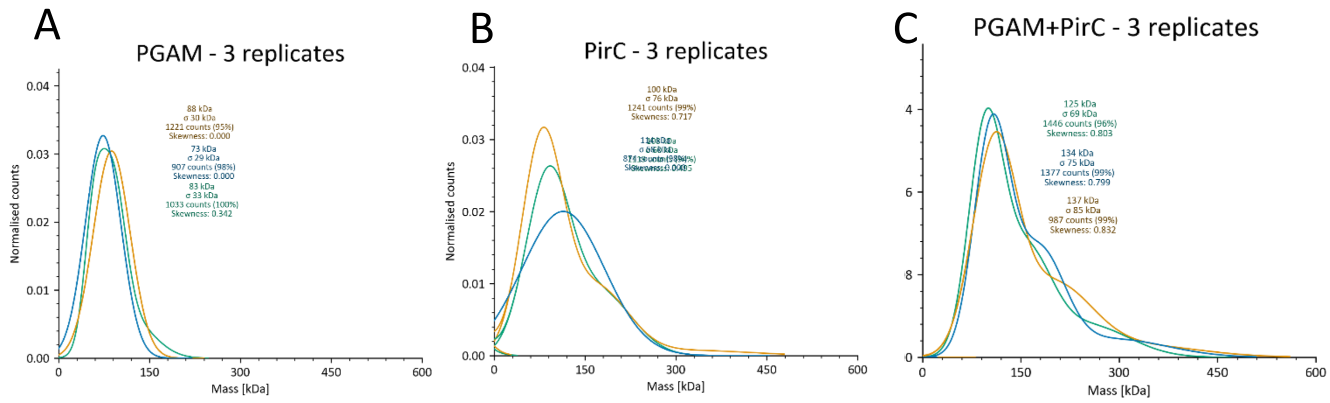


Figure S 3 – Mass photometry of three independent technical replicates of strep-PGAM-WT (A), strep-PirC (B) and the complex of both (C). Gaussian fitting of the counts and the calculated mass of the major mass of the fit. Each bar represents the normalized count.

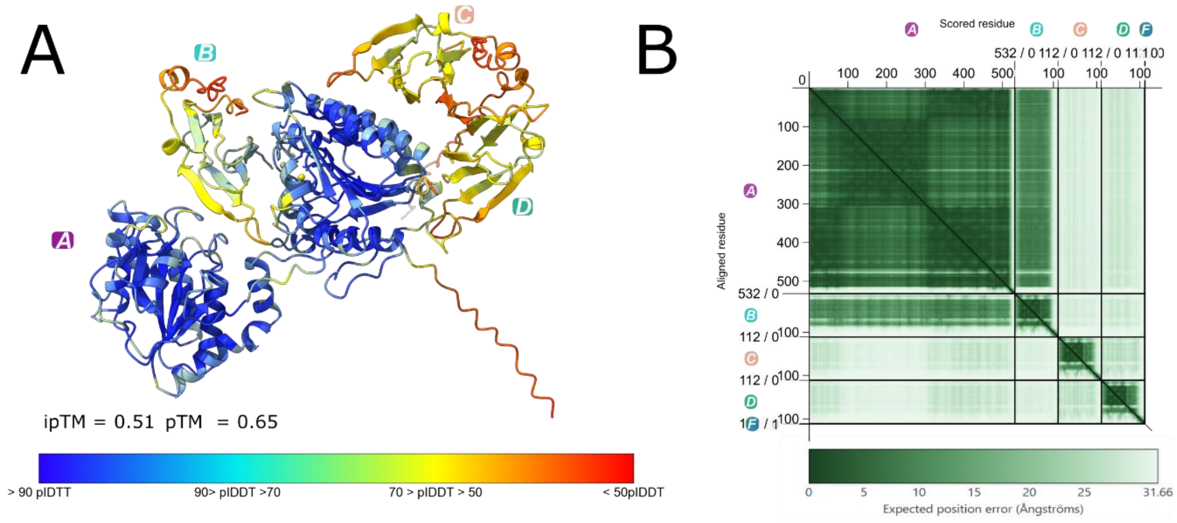


Figure S 4 – Structure of *Synechocystis* PGAM in complex with three monomers of PirC. (A) AlphaFold-predicted structure of the PGAM-PriC(tri) colourised by the predicted IDDT matching score (pIDDT) of the structure (B) Predicted aligned error (PAE) diagram of the PGAM-PriC(tri) complex.

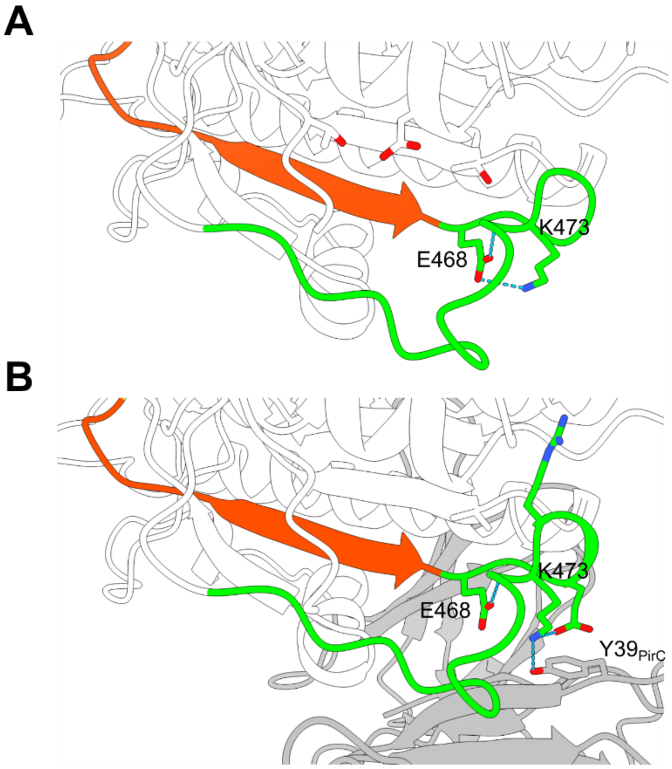


Figure S 5 – Hydrogen bond interactions (A) H-bond of E468 and K473 within the loop (B) H-bond of K473 of the loop with Y39 of PirC.

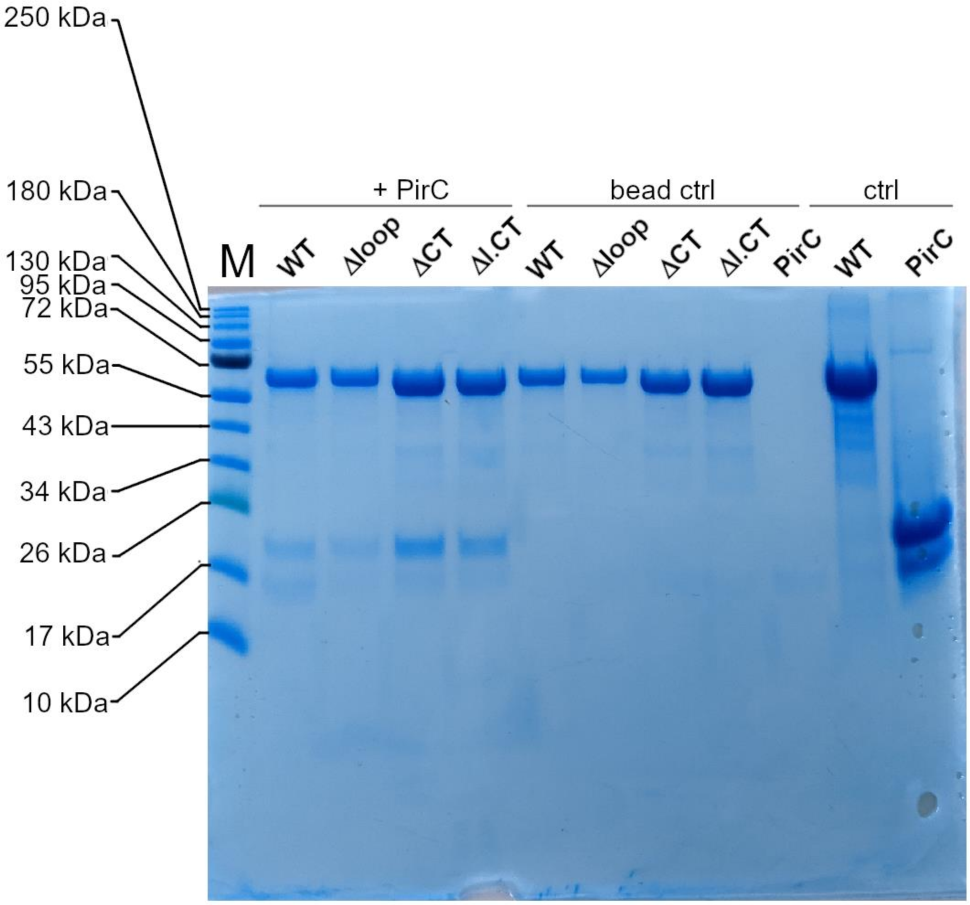


Figure S 6 – Pulldown assay of immobilised strep-PGAM variants on strep-tactin magnetic beads with his-Tagged PirC. The four columns after the standard band M are the samples, the following five columns are the different proteins without interaction incubation. The last two columns are the size control of PGAM and PirC.

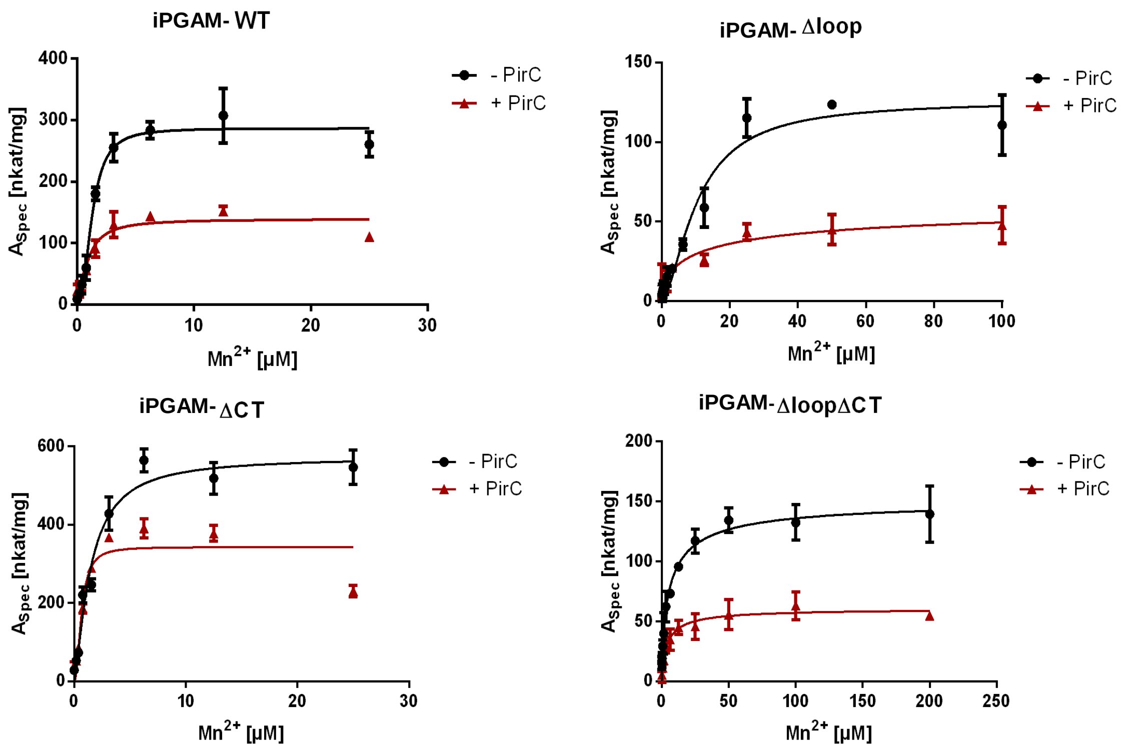


Figure S 7 – Effect of the deletion of the sub-structures on the activity of iPGAM and the inhibition with PirC in dependence of the manganese concentration (A) Hill kinetics of WT without (black) and with PirC inhibition (red) at a concentration of 1.5 mM 3-PGA. Hill coefficiencies: - PirC = 2.319 ± 0.336; + PirC = 1.498 ± 0.333. Khalf: - PirC = 1.268 ± 0.093 µM; +PirC = 0.880 ± 0.150 µM (B) Hill kinetics of $\Delta$loop without (black) and with PirC inhibition (red). Hill coefficiencies: - PirC = 1.605 ± 0.270; + PirC = 0.5798 ± 0,187, Khalf: - PirC = 10.54 ± 1.43 µM; +PirC = 0.1375 ± 18.74µM (C) Hill kinetics of $\Delta$CT without (black) and with PirC inhibition (red). Hill coefficiencies: - PirC = 1.377 ± 0.184; + PirC = 2.079 ± 0.537, Khalf: - PirC = 1.466 ± 0.175 µM; + PirC = 0.666 ± 0.095 µM (D) Hill kinetics of $\Delta$loop$\Delta$CT without (black) and with PirC inhibition (red). Hill coefficiencies: - PirC = 0.728 ± 0.091; + PirC = 0.854 ± 0.181, Khalf: - PirC = 5.781 ± 1.462 µM; +PirC = 4.037 ± 1.300 µM. Each point represents the main of three independent technical replicates. The error bar depicts the standard deviation of the triplicates.

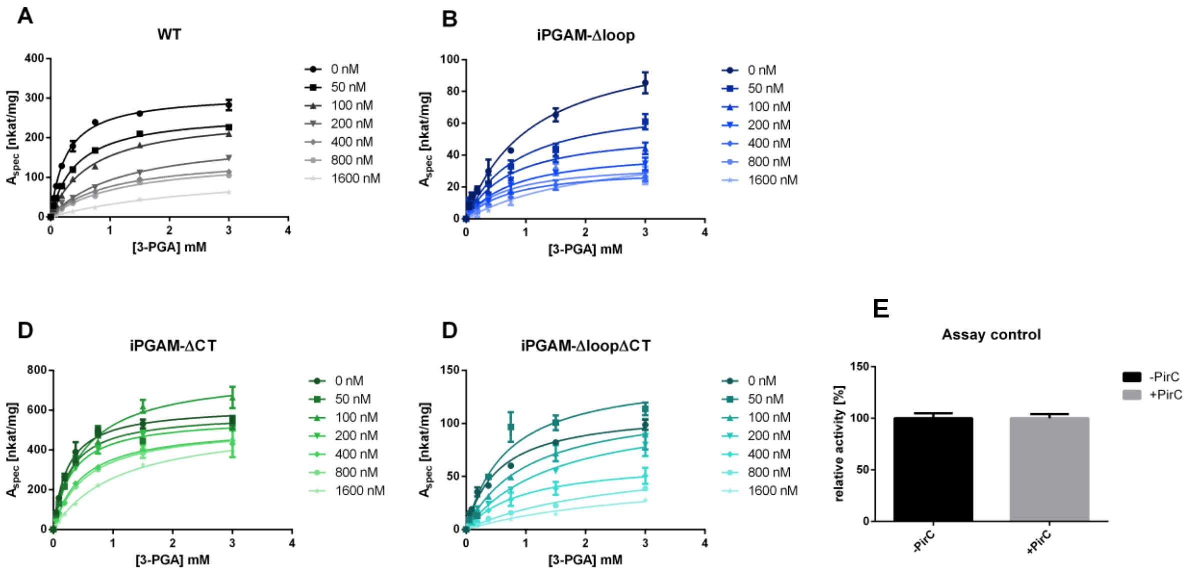


Figure S 8– Effect of PirC on the PGAM variants in varying concentrations of PirC. (A) Michaelis-Menten kinetics of PGAMWT (B) Michaelis-Menten kinetics of PGAM-Δloop. (C) Michaelis-Menten kinetics of PGAM-ΔCT (D) Michaelis-Menten kinetics of PGAMΔloop-ΔCT. Each points represents the main of three independent technical replicates. The error bar depicts the standard deviation of the triplicates. (E) Control of PGAM assay components by adding 2-PGA instead of 3-PGA and 1600 nM PirC.

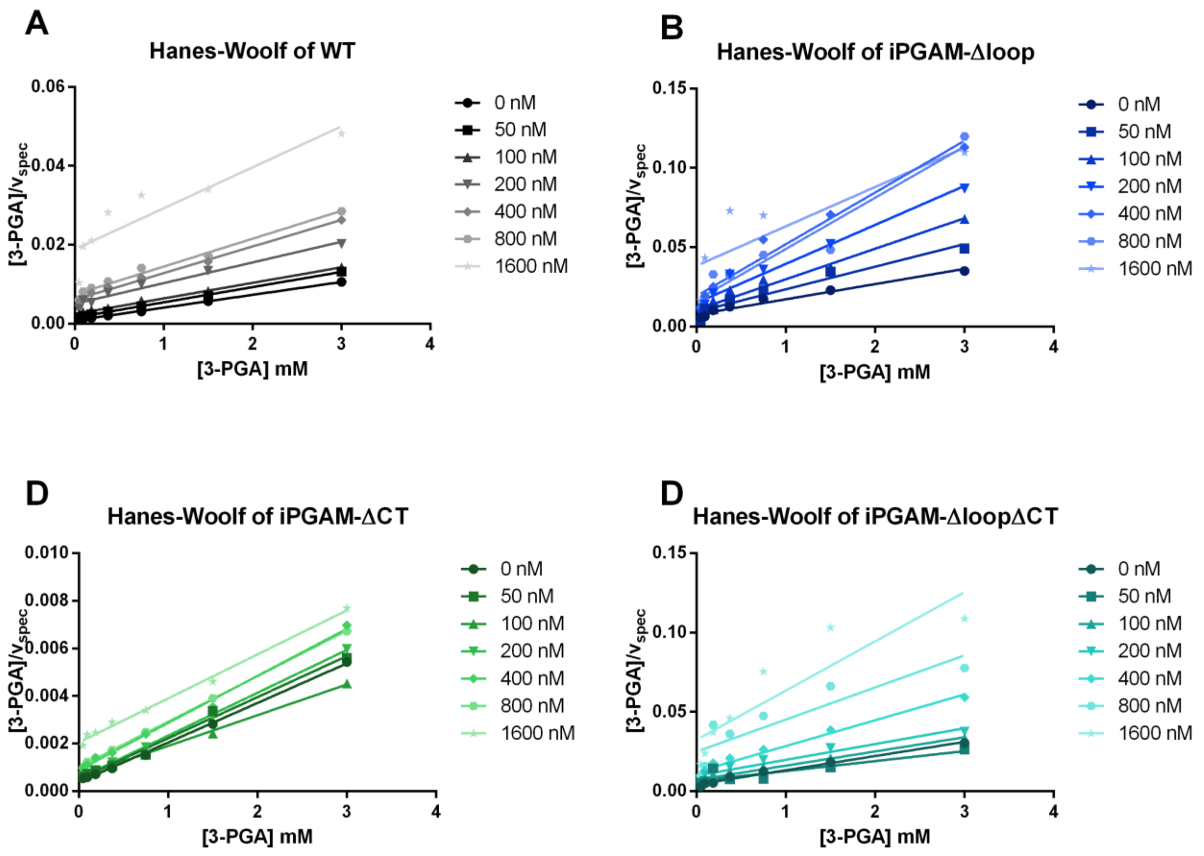


Figure S 9 – Inhibition mechanism and effect of PirC on the PGAM variants in varying concentrations of PirC. (A) Hanes-Woolf kinetics of PGAM-WT(B) Hanes-Woolf kinetics of PGAM-Δloop. (C) Hanes-Woolf kinetics of PGAM-ΔCT (D) Hanes-Woolf kinetics of PGAM-ΔloopΔCT. Each points represents the main of three independent technical replicates. The error bar depicts the standard deviation of the triplicates.

Table S 1 - Calculated kinetic parameters of all iPGAM variants with all concentrations of PirC

iPGAM-WT (50 nM of enzymes used)

| [PirC] [nM] | $K_m$ [mM] | ± [mM] | $v_{max}$ | ± [nkat · $mg^{-1}$] | $k_{cat}$ | ± [$s^{-1}$] | $k_{cat}^{-1} \cdot K_M^{-1}$ | ± [$s^{-1} \cdot M^{-1}$] |
|---|---|---|---|---|---|---|---|---|
| 0 | 0.2648 | 0.01299 | 310.6 | 4.467 | 18.6 | 0.2656 | 70,241.69 | 3,588.78 |
| 50 | 0.4321 | 0.01509 | 263.5 | 3.005 | 15.78 | 0.1822 | 36,519.32 | 1,343.24 |
| 100 | 0.6825 | 0.03328 | 258.5 | 4.828 | 15.51 | 0.2848 | 22,725.27 | 1,184.09 |
| 200 | 1.217 | 0.09253 | 205.7 | 7.031 | 12.33 | 0.4225 | 10,131.47 | 844.93 |
| 400 | 1.018 | 0.07012 | 154.4 | 4.509 | 9.26 | 0.2715 | 9,096.27 | 680.95 |
| 800 | 1.228 | 0.09629 | 150.1 | 5.327 | 8.995 | 0.3189 | 7,324.92 | 630.34 |
| 1600 | 2.895 | 0.3635 | 123.3 | 9.234 | 7.388 | 0.5523 | 2,551.99 | 372.92 |

iPGAM-Δloop

| [PirC] [nM] | $K_m$ [mM] | ± [mM] | $v_{max}$ | ± [nkat · $mg^{-1}$] | $k_{cat}$ | ± [$s^{-1}$] | $k_{cat}^{-1} \cdot K_M^{-1}$ | ± [$s^{-1} \cdot M^{-1}$] |
|---|---|---|---|---|---|---|---|---|
| 0 | 1.087 | 0.1407 | 114.3 | 6.43 | 6.728 | 0.3795 | 6,189.51 | 873.9287499 |
| 50 | 0.8356 | 0.1581 | 73.8 | 5.59 | 4.349 | 0.3297 | 5,204.64 | 1060.852711 |
| 100 | 0.7436 | 0.1396 | 56.11 | 4.069 | 3.301 | 0.2394 | 4,439.21 | 893.4211801 |
| 200 | 0.8023 | 0.1379 | 43.62 | 2.965 | 2.568 | 0.1755 | 3,200.80 | 592.048368 |
| 400 | 0.7501 | 0.1319 | 32.1 | 2.187 | 1.892 | 0.1293 | 2,522.33 | 475.8538227 |
| 800 | 0.665 | 0.1815 | 35.08 | 3.579 | 2.068 | 0.211 | 3,109.77 | 906.1264333 |
| 1600 | 2.288 | 0.7366 | 49.15 | 8.748 | 2.904 | 0.5184 | 1,269.23 | 467.2293431 |

iPGAM-ΔCT

| [PirC] [nM] | $K_m$ [mM] | ± [mM] | $v_{max}$ | ± [nkat · $mg^{-1}$] | $k_{cat}$ | ± [$s^{-1}$] | $k_{cat}^{-1} \cdot K_M^{-1}$ | ± [$s^{-1} \cdot M^{-1}$] |
|---|---|---|---|---|---|---|---|---|
| 0 | 0.2416 | 0.01875 | 617.9 | 13.72 | 35.87 | 0.7918 | 148,468.54 | 11,979.31 |
| 50 | 0.276 | 0.03317 | 583.6 | 20.68 | 33.88 | 1.197 | 122,753.62 | 15,376.95 |
| 100 | 0.5 | 0.03792 | 785.4 | 20.46 | 45.59 | 1.19 | 91,180.00 | 7,313.20 |
| 200 | 0.3083 | 0.01374 | 563.4 | 7.666 | 32.71 | 0.441 | 106,097.96 | 4,940.09 |
| 400 | 0.4819 | 0.056 | 521.5 | 20.66 | 30.27 | 1.199 | 62,813.86 | 7,711.78 |
| 800 | 0.545 | 0.02108 | 525.4 | 7.168 | 30.47 | 0.4157 | 55,908.26 | 2,293.05 |
| 1600 | 1.106 | 0.05293 | 543.9 | 11.38 | 31.55 | 0.6601 | 28,526.22 | 1,489.95 |

iPGAM-ΔloopΔCT

| [PirC] [nM] | $K_m$ [mM] | ± [mM] | $v_{max}$ | ± [nkat · $mg^{-1}$] | $k_{cat}$ | ± [$s^{-1}$] | $k_{cat}^{-1} \cdot K_M^{-1}$ | ± [$s^{-1} \cdot M^{-1}$] |
|---|---|---|---|---|---|---|---|---|
| 0 | 0.5466 | 0.0594 | 112.8 | 4.33 | 6.428 | 0.2468 | 11,759.97 | 1,355.39 |
| 50 | 0.6862 | 0.1473 | 147.2 | 11.89 | 8.418 | 0.6759 | 12,267.56 | 2,811.55 |
| 100 | 1.026 | 0.1573 | 120.3 | 7.874 | 6.856 | 0.4473 | 6,682.26 | 1,113.39 |
| 200 | 1.498 | 0.2363 | 116.5 | 8.865 | 6.648 | 0.5062 | 4,437.92 | 777.34 |
| 400 | 0.9606 | 0.1824 | 65.73 | 5.217 | 3.747 | 0.2968 | 3,900.69 | 802.53 |
| 800 | 2.604 | 0.6414 | 69.9 | 9.917 | 3.974 | 0.5607 | 1,526.11 | 433.20 |
| 1600 | 3.044 | 1.038 | 53.04 | 10.93 | 3.017 | 0.6202 | 991.13 | 394.64 |

### Supplemental Material & Methods

#### Multiple alignments and phylogenetic tree calculation of phosphoglycerate mutases

The alignments were done with Matlab® using the functions `seqdist`, `seqlinkage` and `multialign` in the following script.

```
sequences = table2struct(readtable('data.xlsx')); %loading of the sequences from an excel data and converting to a struct

dist = seqpdist(sequences,'ScoringMatrix','BLOSUM62'); % Calculation of  pairwise distance between sequences based on blosum62 scoring matrix

tree = seqlinkage(dist,'average',sequences); %construction of tree using the pairwise distances

multiple_alignment = multialign(sequences, tree, 'ScoringMatrix', 'BLOSUM80'); %calculation of the multiple alignment using the constructed tree a guide for the progressive alignment and Blosum80 matrix.
```

The resulting tree was visualized with iTol, which was developed by Letunic & Bork (1). It is provided by the EMBL (European Molecular Biology Laboratory) Heidelberg.

#### SWISS-MODEL structure prediction

The structure of the iPGAM was predicted using the SWISS-Model workspace developed by the working group of Thorsten Schwede and published in 2018. SWISS-Model predicts structures by homology modeling against known structures and is provided by the Swiss Institute of Bioinformatics (Lausanne, Switzerland) (2, 3).

### AlphaFold Prediction

The structures were also predicted using the AlphaFoldServer provided by Google (Mountain View, California). It uses the algorithm of AlphaFold 3 developed by Abramson et al. (2024).(5)

### Molecular Cloning, mutagenesis and Plasmids

Gibson Assembly (GA) and mutagenesis PCR were used to create plasmids. The GA was done according to the manufacturer protocol (NEB E2611S/L, E5510S) using a master mix of T5-Exonucleases (0.08 Units), Q5® High-fidelity DNA-Polymerase (0.5 Units) and Taq DNA Ligase (80 Units) (New England Biolabs, Boston) in isobuffer (pH 7.5, 100 mM Tris, 1 mM $MgCl_2$, 1 mM DTT, 0.2 mM dNTP's, 1 mM NAD, and 25% PEG-8000. The assembly fragments were linearized via PCR. To perform the assembly, 5 µl of an assembly mixture was mixed with the master mix and incubated for 60 min at 50 °C.

According to the manufacturer protocol, iPGAM was mutated with the Q5® Site-Directed Mutagenesis Kit (NEB, E0554). Chemical-competent *E. coli* Stellar cells were used for the transformation.

Plasmids created and used in this study are listed in Table S 2.

Table S 2 – List of used plasmids in this study

| Plasmid | Purpose | Source |
|---|---|---|
| pTO104 | Replacement of iPGAM gene (*slr1945*) by mutated iPGAM gene (iPGAM-∆loop) and Spec$^R$ | This study |
| pTO106 | Replacement of iPGAM gene (*slr1945*) by mutated iPGAM gene (iPGAM-∆CT) and Cm$^R$ | This study |
| pTO107 | Replacement of iPGAM gene (*slr1945*) by mutated iPGAM gene (iPGAM-∆loop∆CT) and Cm$^R$ | This study |
| pJA1 | Deletion of *pirC* gene (*sll0944*) via Spec$^R$ cassette | This study |
| pJS15 | Expression of strep-tagged $P_{II}$ protein (Ssl0707) in *E. coli* | (6) |
| pJS22 | Expression of His8-Tagged PirC (Sll0944) in *E. coli* T7-strains | (6) |
| pJS26 | Expression of His8-tagged $P_{II}$ protein (Ssl0707) in *E. coli* T7-strains | (6) |
| pJS27 | Expression of strep-tagged PirC (Sll0944) in *E. coli* | (7) |
| pET-28a(+)-iPGAM | Expression His8-tagged-iPGAM (Slr1945) in *E. coli* T7-strains | (7) |
| pTO301 | Expression of His8-tagged iPGAM-∆loop in *E. coli* | This study |
| pTO304 | Expression of strep-tagged iPGAM in *E. coli* | This study |
| pTO305 | Expression of strep-tagged iPGAM-∆loop in *E. coli* | This study |
| pTO307 | Expression of strep-tagged iPGAM-∆CT in *E. coli* | This study |
| pTO308 | Expression of strep-tagged iPGAM-∆loop∆CT in *E. coli* | This study |

### Strains

The strains used in this study are listed in Table S 3

Table S 3 – List of used organisms in this study

| Organism | Strain | Genotype | Purpose |
|---|---|---|---|
| *E. coli* | NEB10β | *Δ(ara-leu) 7697 araD139 fhuA ΔlacX74 galK16 galE15 e14- Φ80dlacZΔM15 recA1 relA1 endA1 nupG rpsL (StrR) rph spoT1 Δ(mrr-hsdRMS-mcrBC)* | Molecular Cloning |
| *E. coli* | Lemo21(DE3) | *fhuA2 [lon] ompT gal (λ DE3) [dcm] ΔhsdS/ pLemo(CamR)*<br>*λ DE3 = λ sBamHIo ΔEcoRI-B int::(lacI::PlacUV5::T7 gene1) i21 Δnin5*<br>pLemo = pACYC184-*PrhaBAD-lysY* | Protein Expression |
| *Synechocystis* sp. PCC 6803 | wild type glucose-sensitive | WT | Background strain, control |
| *Synechocystis* sp. PCC 6803 | ΔPirC | *sll0944*::Spec$^{R}$ | Control strain (This study) |
| *Synechocystis* sp. PCC 6803 | iPGAM-Δloop | *slr1945*::PGAMΔloop-Spec$^{R}$ | iPGAM variant testing (This study) |
| *Synechocystis* sp. PCC 6803 | iPGAM-ΔCT | *slr1945*::iPGAMΔCT-Cm$^{R}$ | iPGAM variant testing (This study) |
| *Synechocystis* sp. PCC 6803 | iPGAM-ΔloopΔCT | *slr1945*::iPGAMΔloop ΔCT-Spec$^{R}$ | iPGAM variant testing (This study) |

**Cultivation of Cyanobacteria**

Growth experiments and precultures of *Synechocystis* were cultivated in $BG_{11}$ with the composition explained by (34). Baffle-free 200 ml Erlenmeyer flasks were used for 50 ml cultures. Standard cultivation was performed at 28 °C with continuous shaking at 125 rpm, either at constant (24 h/d, ~50 µE · $m^{-2}$ · $s^{-1}$) or fluctuating (12 h light;~50 µE · $m^{-2}$ · $s^{-1}$/12 h dark) illumination. For different experiments, the BG11 was adjusted as listed in Table S 4. When necessary, appropriate antibiotics were supplemented in the media to ensure the continuity of the mutation. The growth was recorded by measuring the $OD_{750}$.

Table S 4– Exchanges in media composition for different purposes

| **Type of Experiment** | **Adjustment** |
|---|---|
| Control | no adjustment |
| Ammonium growth | Replacement of $NaNO_4$ to 5 mM Ammonium chloride. The lost $Na^+$ ions were re-supplemented by 17 mM NaCl. ($BG_{11,\ Ammonia}$) |
| Nitrogen depletion | Removal of $NaNO_4$ and supplementation of 17 mM NaCl. $BG_{11,0}$ |

For nitrogen deficiency experiments, precultures of *Synechocystis* were cultivated for three days, as described previously, at an initial $OD_{750}$ of 0.1. Experimental cultures were prepared in BG11 medium with a set starting $OD_{750}$ of 0.2 and grown for two days under identical conditions until they reached an $OD_{750}$ of 0.6-1. For the nitrogen depletion experiments, cells from the cultures were harvested by centrifugation (4000 g, 10 min), washed with and resuspended in $BG_{11,0}$ medium to create cultures with an initial $OD_{750}$ of 0.4.

For ammonium experiments, cells were diluted in fresh $BG_{11}$, $_{\text{Ammonia}}$ medium and adjusted to an $OD_{750}$ of 0.2.

*Escherichia coli* cultures were grown in LB medium and on LB agar. Lennox broth: 5 $g \cdot l^{-1}$ Yeast extract, 10 $g \cdot l^{-1}$ Tryptone, NaCl 5 $g \cdot l^{-1}$, and solid: 15 $g \cdot l^{-1}$ agar were used.

**Expression and Purification of Proteins**

*E. coli* Lemo21(DE3) was used for the overexpression of the various kinds of proteins. His-tagged proteins were expressed as described in the manufactured expression protocol in 2-fold concentrated LB media with appropriate antibiotics. An overnight expression was induced by adding 400 µM IPTG at 20 °C during continuous shaking at 120 rpm. The expression of strep-tagged proteins based on pASK-Iba5Plus expression plasmid was induced by adding 200 $\mu g \cdot l^{-1}$ anhydrotetracycline.

The heterologous proteins containing His-tags were purified via 5 ml Ni-NTA HisTrap columns (Cytiva, Marlborough, USA). The cells were lysed in 50 ml lysis buffer containing 50 mM Tris/HCl buffer pH 8.0, 300 ml NaCl, 1 mM DTT, and cOmplete™ (Roche, Basel, Switzerland). The His-tagged proteins were loaded on the Ni-NTA column with Buffer A containing 50 mM Tris/HCl pH 8.0, 300 ml NaCl and eluted via a gradient of increasing imidazole (0-500 mM,

Buffer B) using a ÄKTAPurifier™ System (Cytiva, Marlborough, USA). After this first purification, the proteins were purified further via size exclusion chromatography using a Superdex™ 200 Increase 10/300 GL (Cytiva, Marlborough, USA) with 50 mM Tris/HCl buffer containing 100 mM KCl and 0.5 mM EDTA.

5 ml Strep-Tactin® Superflow® columns (IBA Lifescience, Göttingen, Germany) were used to purify Strep-tagged proteins. Cells were lysed in lysis buffer containing 100 mM Tris/HCl pH 8.0, 150 mM NaCl, and cOmplete™ (Roche, Basel, Switzerland). The proteins were loaded on the column and eluted with a buffer containing 2.5 mM Desthiobiotin. The buffer of each purified protein was exchanged via dialysis using dialysis buffer (50 mM Tris/HCl pH 8.0, 100 mm KCl, 40% glycerol) and a 3.5 kDa cutoff dialysis tube. According to previous studies, all purification steps were checked via SDS-PAGE.

**Mass photometry using the Refyn OneMP**

A mass photometry experiment was used to study the variants' oligomerization and the stoichiometry of the iPGAM-PirC complex. The measurement was done according to Wu & Piszczek (9). Allexperiments were performed in a buffer containing 25 mM Tris-HCl, 50 mM KCl and 50 µM $MnCl_2$ at pH 8.0. The buffer and all samples were filtered through a 0.22 µM sterile filter, and the buffer was degassed by vacuum degassing. As coverslips, the High-Precision Microscope Cover Glasses (Marienfeld, Lauda-Königshofen, Germany) were used, which were cleaned three times in ultrapure water and 100% isopropanol under a sterile hood. The coverslips were adjusted onto the objective, and the CultureWell™ Reusable Gasket was placed on it. The machine was focused by adding 10 µl of the buffer in the gaskets on the coverslips. Afterwards, 10 µl of the sample was added to the buffer and measured for 1 min. The Refyn OneMP was calibrated using ovalbumin (43 kDa, Hen egg), cobalamin (75 kDa, chicken egg white), and aldolase (157 kDa, rabbit muscle) from the Gel Filtration Calibration Kit LMW (Cytiva, Marlborough, USA). The data was analyzed using the DiscoverMP® software.

**BLI using the Octet K2 System**

*In vitro* binding studies were done using bio-layer interferometry (BLI) using the Octet K2 system (Sartorius, Göttingen, Germany) according to the Bio-protocol (10). The experiments were performed in HEPES buffer (20 mM HEPES-KOH pH 8.0, 50 µM $MnCl_2$, 0.005 % NP-40, In the first step, His6-iPGAM (1000 nM) were immobilized on Ni-NTA sensors (Satorius), followed by a 60 sec baseline measurement. For the binding of PirC, the biosensors were dipped into the PirC solution for 180-sec (Association), with concentrations ranging between 9.375 nM – 1500 nM. A 300-sec dissociation step terminated the assay. One measurement without any interaction partner was performed to prevent false positive results in each experimental set. The biosensors were regenerated after each use with 10 mM glycine (pH 1.7) and 10 mM $NiCl_2$, as proposed in manufacturer recommendations. The recorded curves of a set were preprocessed by aligning them to the average of the baseline step and the dissociation step. The response in equilibrium (Req) was calculated using the Data Analysis Software of the Octet System. The concentration versus Req plots were made.

**Phosphoglycerate Mutase Assay**

The iPGAM activity was determined by a coupled enzyme assay as adapted as described previously (7). Around 0.6 µg of purified iPGAM was used in a 200 µl reaction. The reaction mixture containing 20 mM HEPES-KOH (pH 8.0), 100 mM KCl, 5 mM $MgSO_4$, 0.2 mM $MnCl_2$, 50 µg · $ml^{-1}$ BSA, 1 mM DTT, 0.4 mM ADP, 0.2 mM NADH, 0.1 U enolase (Sigma Aldrich, St. Louis, USA), 0.4 U pyruvate kinase (Roche, Basel, Switzerland), 0.4 U lactate dehydrogenase (Roche, Basel, Switzerland) and 1 µg iPGAM was pre-warmed to 30 °C. The assay was started by adding the 3-PGA solutions. The resulting decrease of NADH over time was recorded in a 96-well plate using BioTek Epoch 2 (Agilent, Santa Clara) at 340 nm. A blank assay without 3-PGA was also performed, and no decrease was detectable.

**Glycogen Measurement**

The glycogen content was quantified according to previous studies from 2 ml *Synechocystis* culture samples (11). The glycogen was hydrolyzed to glucose with 4.4 U · $µl^{-1}$ amyloglucosidase from *Aspergillus niger* (Sigma Aldrich, St. Louis, USA) for 2 h at 60 °C. The resulting glucose concentration was measured via an *o*-toluidine assay (12). The samples were boiled in 1:6

dilution with a 6% *o*-toluidine reagent (in glacial acetic acid) for 10 min, then cooled on ice and measured with Spark M10 (Tecan, Männedorf, Switzerland) at 635 nm. The concentration of samples was calculated using a calibration curve of a defined quantity of glucose (0, 10 µg, 50 µg, 100 µg, 250 µg, and 500 µg).

**PHB Quantification**

As described previously, polyhydroxybutyrate (PHB) was detected using high-performance liquid chromatography (HPLC) (7, 13, 14).

### References

1. I. Letunic, P. Bork, Interactive Tree of Life (iTOL) v6: recent updates to the phylogenetic tree display and annotation tool. *Nucleic Acids Res* 52, W78–W82 (2024).

2. N. Guex, M. C. Peitsch, T. Schwede, Automated comparative protein structure modeling with SWISS-MODEL and Swiss-PdbViewer: A historical perspective. *Electrophoresis* 30, S162–S173 (2009).

3. A. Waterhouse, *et al.*, SWISS-MODEL: homology modelling of protein structures and complexes. *Nucleic Acids Res* 46, W296 (2018).

4. J. Abramson, *et al.*, Accurate structure prediction of biomolecular interactions with AlphaFold 3. *Nature 2024 630:8016* 630, 493–500 (2024).

5. J. Abramson, *et al.*, Accurate structure prediction of biomolecular interactions with AlphaFold 3. *Nature 2024 630:8016* 630, 493–500 (2024).

6. J. Scholl, L. Dengler, L. Bader, K. Forchhammer, Phosphoenolpyruvate carboxylase from the cyanobacterium Synechocystis sp. PCC 6803 is under global metabolic control by PII signaling. *Mol Microbiol* 114, 292–307 (2020).

7. T. Orthwein, *et al.*, The novel PII-interactor PirC identifies phosphoglycerate mutase as key control point of carbon storage

metabolism in cyanobacteria. *Proc Natl Acad Sci U S A* 118, e2019988118 (2021).

8. M. Mager, *et al.*, Interlaboratory Reproducibility in Growth and Reporter Expression in the Cyanobacterium Synechocystis sp. PCC 6803. *ACS Synth Biol* 12, 1823–1835 (2023).

9. D. Wu, G. Piszczek, Standard protocol for mass photometry experiments. *European Biophysics Journal* 50, 403–409 (2021).

10. T. Orthwein, L. F. Huergo, K. Forchhammer, K. A. Selim, Kinetic analysis of a protein-protein complex to determine its dissociation constant (kd) and the effective concentration (ec50) of an interplaying effector molecule using bio-layer interferometry. *Bio Protoc* 11 (2021).

11. S. Doello, A. Klotz, A. Makowka, K. Gutekunst, K. Forchhammer, A Specific Glycogen Mobilization Strategy Enables Rapid Awakening of Dormant Cyanobacteria from Chlorosis. *Plant Physiol* 177, 594–603 (2018).

12. K. M. DUBOWSKI, An o-Toluidine Method for Body-Fluid Glucose Determination. *Clin Chem* 8, 215–235 (1962).

13. M. Koch, S. Doello, K. Gutekunst, K. Forchhammer, PHB is Produced from Glycogen Turn-over during Nitrogen Starvation in Synechocystis sp. PCC 6803. *Int J Mol Sci* 20, 1942 (2019).

14. M. Koch, *et al.*, Maximizing PHB content in Synechocystis sp. PCC 6803: a new metabolic engineering strategy based on the regulator PirC. *Microb Cell Fact* 19, 1–12 (2020).
